# CRIC-seq reveals positional rule of PTBP1-mediated long-range RNA looping in splicing regulation

**DOI:** 10.1101/2022.08.09.503273

**Authors:** Rong Ye, Naijing Hu, Changchang Cao, Ruibao Su, Chen Yang, Shihan Xu, Yuanchao Xue

**Affiliations:** Key Laboratory of RNA Biology, Institute of Biophysics, Chinese Academy of Sciences, Beijing 100101, China; University of Chinese Academy of Sciences, Beijing 100049, China; State Key Laboratory of Ophthalmology, Optometry and Vision Science, Wenzhou, Zhejiang, 325003, China

**Author notes:** Contributed equally (Rong Ye, Naijing Hu).

**Keywords:** RNA-binding protein, RNA-RNA interaction, Alternative splicing, PTBP1, Deep sequencing

## Abstract

RNA-binding proteins bind at different positions of pre-mRNA molecules to promote or reduce the usage of a particular exon. Seeking to understand the working principle of these positional effects, we develop a CRIC-seq method to enrich single RBP-mediated in situ RNA-RNA spatial interacting fragments for deep sequencing. We determine hnRNPA1- and PTBP1-mediated RNA-RNA interactions and regulatory mechanisms in HeLa cells. Unexpectedly, 3D RNA map analysis shows that PTBP1-mediated loops in introns preferably promote cassette exon splicing by accelerating asymmetric intron removal, whereas the loops spanning across cassette exon primarily repress splicing. This “positional rule” can faithfully predict PTBP1-regulated splicing outcomes. We further demonstrate that cancer-related splicing quantitative trait loci can disrupt RNA loops by reducing PTBP1 binding on pre-mRNAs to cause aberrant splicing in tumors. Our study presents a powerful method for exploring the functions of RBP-mediated RNA-RNA interactions in gene regulation and disease.

## INTRODUCTION

Eukaryotic organisms use alternative splicing to generate multiple different isoforms from a single precursor mRNA (pre-mRNA) to execute similar or completely different molecular functions (Baralle and Giudice, 2017; Black, 2003; Bonnal et al., 2020; Gabut et al., 2011). Alternative splicing is carried out by spliceosome, which can faithfully recognize splice signals to remove the much longer intronic sequences and simultaneously join the shorter alternative exons (Black, 2003). In addition to canonical splice signals, various auxiliary sequences within introns and exons can be bound by different RNA-binding proteins (RBPs) such as Nova, PTBP1, and hnRNPA1 to promote or repress splicing (Fairbrother et al., 2002; Fu and Ares, 2014; Sharma et al., 2008; Ule et al., 2006; Van Nostrand et al., 2020; Wang et al., 2004). Global mapping of protein-RNA interaction sites by CLIP methodologies has revealed an emerging theme: RBP binding at different positions tends to cause different splicing outcomes (Ule et al., 2006; Van Nostrand et al., 2020; Xue et al., 2009; Yeo et al., 2009). As a long-standing puzzle, how the positional effects are achieved and what are the underlying mechanisms remain unclear.

PTBP1 was initially characterized as a splicing repressor and prefers to bind CU-rich motifs (Singh et al., 1995; Spellman and Smith, 2006). In addition to repressed exons, previous microarray analyses had identified many PTBP1-activated exons (Boutz et al., 2007; Llorian et al., 2010; Xing et al., 2008). Aligning the 1D landscape of PTBP1 binding with the splicing outcome further uncovered a general trend: prevalent PTBP1 binding near the splice sites of cassette exon represses splicing (Van Nostrand et al., 2020; Xue et al., 2009), whereas the dominate binding near constitutive splice sites promotes splicing (Xue et al., 2009). Like hnRNPA1 and hnRNPK (Blanchette and Chabot, 1999; Cai et al., 2020), PTBP1 also functions as a homodimer within the nucleus (Xue et al., 2009). The dimerization and oligomerization of PTBP1 might induce RNA looping to sequester cassette exons from the splicing machinery or form a zone to block the recognition of splice sites (Black, 2003). Additionally, PTBP1 contains four RRMs, which may individually bind CU-rich RNA elements at different locations to induce RNA looping (Oberstrass et al., 2005). Based on these findings, two hypothetical “loop out of exon” and “loop out of branch point” models have been proposed to explain PTBP1-mediated splicing repression (Oberstrass et al., 2005; Spellman and Smith, 2006). As direct experimental evidence is lacking, these models’ validity still needs to be examined.

RBPs co-transcriptionally bind to nascent pre-mRNA transcripts to facilitate their folding into intricate structures for subsequent processing (Herzel et al., 2017; Xue, 2022). CLIP methodologies could identify the 1D landscape of RBP binding to pre-mRNAs (Lee and Ule, 2018). However, the spatial interactions between these individual binding sites are unknown. Several state-of-the-art methods such as CLASH, hiCLIP, and irCLASH have been developed to profile single protein-mediated RNA-RNA interactions (Helwak et al., 2013; Kudla et al., 2011; Song et al., 2020; Sugimoto et al., 2015). However, the ectopic expression of a defined RBP in cells may disturb the native RNA regulatory networks, and the proximity ligation (in a diluted solution) used by these methods is prone to introduce many false-positive chimeric interactions (Xue, 2022). The limitations restrict the application of these methods for studying the role of RNA-RNA interactions in splicing regulation.

RBPs can bind, stabilize or destabilize RNA-RNA interactions (Cai et al., 2020; Dominguez et al., 2018; Honig et al., 2002; Warf et al., 2009). RNA-RNA interactions around canonical splice signals have been proposed to repress exon usage by blocking 5’ or 3’ splice sites (SS) and the branch point sequence recognition (Blanchette and Chabot, 1997; Liu et al., 2002; Singh et al., 2007; Varani et al., 1999; Warf and Berglund, 2010; Watakabe et al., 1989). Conversely, long-range RNA loops away from the splice signals may promote exon usage by bringing the 5’ SS and 3’ SS into proximity (Charpentier and Rosbash, 1996; Goguel and Rosbash, 1993; Howe and Ares, 1997; Meyer et al., 2011; Singh et al., 2007; Xu et al., 2021), and such hypothetical interactions even enable the bypass of the requirement for a core splicing component U2AF2 in some zebrafish genes (Lin et al., 2016). Moreover, a recent global mapping of nascent RNA structure in HEK293 cells reveals an unexpected trend that the co-transcriptionally well-spliced introns are more structured than poorly spliced ones (Saldi et al., 2021). These studies underscore the critical role of RNA-RNA interactions in splicing regulation.

We previously developed RIC-seq for global profiling of all the RBP-mediated RNA-RNA spatial interactions in an unbiased manner (Cai et al., 2020). However, which RBP mediates the identified RNA proximal interactions is unknown. To overcome this and the aforementioned limitations, here we developed a captured RIC-seq (CRIC-seq) method to profile the RNA interactome arranged by a single RBP. Using CRIC-seq method, we successfully captured and validated RNA-RNA interactions organized by PTBP1 and hnRNPA1 in HeLa cells. Such unbiased mapping enabled us to construct a 3D RNA map to predict PTBP1-mediated RNA loops in promoting or repressing cassette exon splicing in a position-dependent manner. We also demonstrated that cancer-related splicing quantitative trait loci (sQTLs) could disrupt PTBP1-mediated loops to induce tumorigenesis. Together, our results show that CRIC-seq is a powerful method to profile single protein-mediated RNA-RNA spatial interactions in transcriptional and post-transcriptional regulations.

## RESULTS

### Overview of CRIC-seq method

Seeking to profile a single RBP-mediated RNA-RNA interactions at high resolution, we developed the “CRIC-seq” method that leverages the principle of RIC-seq and immunoprecipitation (IP)-mediated enrichment strategy. Briefly, around 1 × 10^7^ cells (*e.g.*, HeLa) are cross-linked with formaldehyde to fix all of the protein-mediated RNA-RNA interactions in living cells (Figure 1A). After permeabilization and micrococcal nuclease treatment, the RNA fragments in close proximity are labeled with pCp-biotin and then ligated *in situ* using T4 RNA ligase. The cells are lysed by brief sonication and then incubated with a specific antibody (for the user-defined analyte RBP) to pull down the RBP-organized whole complement of RNA-RNA spatial interactions (Figure 1A). After RNA extraction and fragmentation, the chimeric RNAs with C-biotin at the juncture of two interacting fragments are enriched using streptavidin magnetic beads for library construction and subsequent paired-end sequencing (Figure 1A). To exclude non-specific background interactions during two sequential IP steps, we used two negative controls for all CRIC-seq experiments: IgG IP with pCp-biotin labeling, and specific antibody IP but without pCp-biotin labeling.

**Figure 1.**
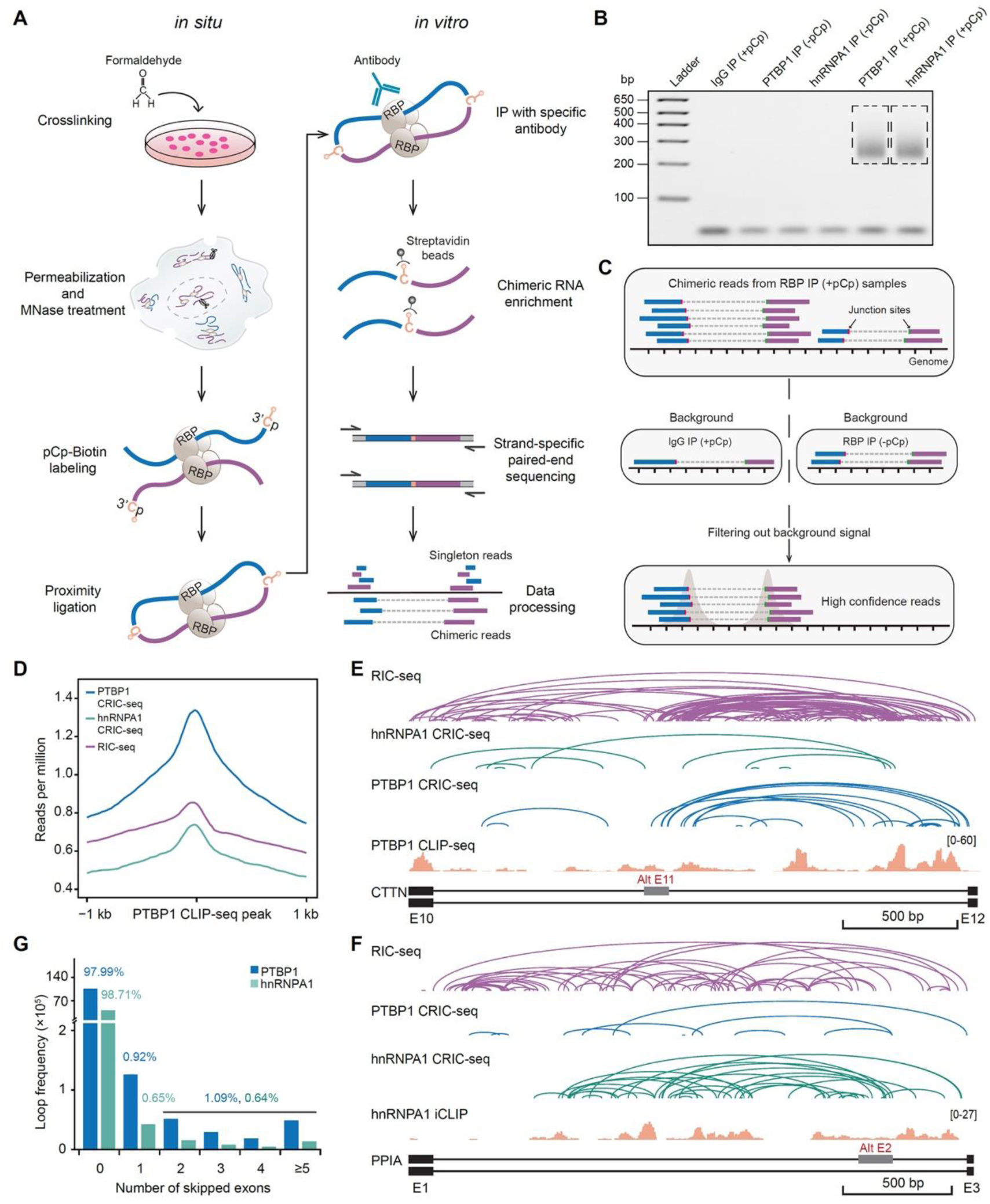
CRIC-seq captures RBP-mediated RNA spatial interactions in HeLa cells. (A) Diagram of the CRIC-seq method. Single RBP-mediated proximal interacting RNAs were first fragmented and labeled with pCp-biotin for in situ proximity ligation. The ligated RNAs were sequentially selected with a specific antibody and streptavidin beads by immunoprecipitation. The captured RNAs with an additional C at the juncture were subsequently converted into paired-end libraries for deep sequencing. MNase, micrococcal nuclease. (B) Agarose gel showing the size distribution of PTBP1 and hnRNPA1 CRIC-seq libraries. (C) Schematic diagram of CRIC-seq data analysis pipelines. (D) Meta gene profile showing PTBP1 CRIC-seq (blue), hnRNPA1 CRIC-seq (green), and RIC-seq (purple) signals around the PTBP1 CLIP-seq peaks identified in HeLa cells. (E, F) Snapshot of CRIC-seq, RIC-seq, and CLIP-seq signals around *CTTN* exon 11 and *PPIA* exon 2. Representative tracks showing high confidence RIC-seq loops (purple arc lines), PTBP1 CRIC-seq loops (blue arc lines), hnRNPA1 CRIC-seq loops (green), and CLIP-seq (orange) signals. (G) Bar graph showing the number of skipped exons for PTBP1 or hnRNPA1-mediated intramolecular RNA loops that located at introns. The percentage of each category is indicated.

### CRIC-seq identifies RBP-mediated RNA spatial interactions

We next chose PTBP1 and hnRNPA1 for CRIC-seq library construction due to their well-known RNA binding motifs and targets (Bruun et al., 2016; Burd and Dreyfuss, 1994; Haberman et al., 2017; Huelga et al., 2012; Oberstrass et al., 2005; Xue et al., 2009). We prepared two biological replicates of CRIC-seq libraries for the PTBP1 IP (+pCp) and hnRNPA1 IP (+pCp) samples using HeLa cell lysates. Three negative controls were also included: IgG IP (+pCp), PTBP1 IP (−pCp), and hnRNPA1 IP (−pCp) (Figure 1B). In contrast to these three negative controls and no template control, we observed a smeared band between 225 bp and 450 bp, with the highest concentration at 271 bp and 288 bp, for the PTBP1 and hnRNPA1 IP (+pCp) samples (Figures 1B and S1A). After purifying the bands with similar size ranges in different libraries for deep sequencing, we obtained approximately 0.15 million, 0.07 million, 0.01 million, 57.95 million, and 26.79 million chimeric reads for PTBP1 IP (−pCp), hnRNPA1 IP (−pCp), IgG IP (+pCp), PTBP1 IP (+pCp), and hnRNPA1 IP (+pCp) samples, respectively (Figure S1B). The low sequencing reads obtained from control samples highlight the specificity of pCp-biotin labeling and antibody IP steps in our CRIC-seq method.

CRIC-seq libraries for the two biological replicates of PTBP1 (+pCp, *R*=0.96) and hnRNPA1 (+pCp, *R*=0.95) were highly reproducible (Figure S1C). In contrast, the *Pearson’s* correlation coefficients for IgG (+pCp) and PTBP1 (−pCp) negative controls were as low as 0.10 and 0.03, respectively. As the chimeric reads from two negative controls represented non-specific background interactions, we thus merged two replicates of each sample for subsequent analyses (Figure 1C). Aiming to enable confident identification of authentic RNA-RNA interactions mediated by PTBP1 or hnRNPA1, we developed a data analysis pipeline: briefly, RNA chimeras from IgG (+pCp) and RBP IP (−pCp) samples were first combined to define the background level. Then, the chimeric reads from the RBP (+pCp) samples were retained based on whether the read number was more than 5-fold greater than the background value defined for the same locus. Finally, the remaining RBP IP (+pCp) chimeric reads with more than two overlapping junctions within 10-nt window sites were assembled into individual clusters. We obtained 4,702,162 clusters supported by 17,966,354 chimeric reads for PTBP1, and 2,185,516 clusters supported by 8,517,935 chimeric reads for hnRNPA1 (Figure 1C and Table S1).

We next compared the single RBP-mediated RNA spatial interactions identified from CRIC-seq with two previously published datasets for HeLa cells: a RIC-seq study of the RNA interactome mediated by all RBPs and the 1D RBP binding sites revealed by a CLIP-seq analysis. We found that 26% of the PTBP1 and 35% of the hnRNPA1 CRIC-seq-identified clusters were among the RNA-RNA interaction clusters revealed by RIC-seq (Figure S1D), indicating that these RIC-seq clusters were mediated by PTBP1 and hnRNPA1. In addition, PTBP1 CRIC-seq signals peaked at the CLIP-seq-identified PTBP1 binding sites, and showed more than 1.6-fold of signal strength than RIC-seq and hnRNPA1 CRIC-seq (Figure 1D), highlighting the power of our method to enrich specific RBP-mediated RNA-RNA interactions. Similar to CLIP-seq, CRIC-seq recapitulated dominant PTBP1 binding on the distal constitutive 3’ splice site of *CTTN* cassette exon 11 (Xue et al., 2009). Moreover, we identified multiple long-range RNA-RNA interactions between this distal constitutive 3’ splice site and the proximal 5’ splice site of *CTTN* exon 11 (Figure 1E), and such PTBP1-mediated looping patterns were not observed for hnRNPA1. On the contrary, hnRNPA1-regulated splicing of *PPIA* exon 2 possesses dominant hnRNPA1 but sparse PTBP1 CRIC-seq signals (Figure 1F).

We noted that PTBP1 and hnRNPA1 CRIC-seq reads (singleton and chimeric reads) were highly correlated with their CLIP-seq reads (*R*=0.89 or 0.86, Figure S1E), suggesting that CRIC-seq detected both 1D RBP binding sites and their 3D interactive interactions. Although over 80% of the PTBP1 CRIC-seq chimeric reads overlapped with CLIP-seq reads (see Type II and III, Figure S1F), we noted 3,075,083 CRIC-seq reads (Type I) did not have the CLIP-seq binding evidence. This might be due to the higher sequencing depth of PTBP1 CRIC-seq over CLIP-seq (57.9 million vs. 31.4 million). Motif analysis of these CRIC-seq chimeric sequences revealed several CU-rich and poly(U) motifs, indicating that these positions might also be bound by PTBP1 or its associated RBPs revealed by STRING database (Figures S1G and S1H). Collectively, these results demonstrate that CRIC-seq can faithfully identify single RBP-mediated RNA spatial interactions.

### Features of PTBP1-mediated RNA spatial interactions

Since PTBP1 is emerging as a master regulator of cellular reprogramming and tumorigenesis (Hu et al., 2018), we thus focus on PTBP1-mediated RNA-RNA spatial interactions to investigate which processes they were most likely involved. We noted that approximately 90% of the PTBP1 CRIC-seq chimeric reads and 96% of the hnRNPA1 CRIC-seq chimeric reads were formed between different fragments in the same RNA molecules (Figure S1I), indicating that these two RBPs may predominantly mediate intramolecular RNA-RNA interactions within cells. For PTBP1 intramolecular reads, we found that their spanning distances ranged from several to millions of nucleotides, with 58.4% shorter than 10 nt, and 28% longer than 100 nt (Figure S1J). A similar trend was also observed for hnRNPA1-mediated RNA-RNA interactions. In addition, genomic distribution analysis showed that 85% of the PTBP1 chimeric reads were located at introns, with 2% mapped to exons, and 6% resided in 3’UTRs (Figure S1K, left panel). For comparison, hnRNPA1 intramolecular chimeric reads showed a similar distribution pattern (Figure S1K, right panel).

RBP-mediated intramolecular RNA-RNA interactions in introns can drive the formation of various RNA loops (Xue, 2022). We next examined the feature of PTBP1-mediated looping patterns. We found that 1.1% of the PTBP1-mediated intronic RNA loops could span multiple exons, and approximately 98% of the loops were restricted within introns, which could not span across any exons (Figure 1G). As a control, hnRNPA1-mediated intramolecular RNA-RNA interactions showed nearly identical looping patterns. Moreover, meta gene analysis revealed that the PTBP1 and hnRNPA1 intramolecular RNA loops preferably happened within the same introns and peaked around intron centers (Figure S1L). These binding and looping features suggest functional implications of PTBP1-mediated RNA loops in splicing regulation, which agree well with its known molecular function.

### PTBP1-mediated RNA loops in splicing regulation

To explore the potential impacts of PTBP1-mediated RNA loops in regulated splicing, we used rMATS to identify putative PTBP1-regulated splicing events in a previously published RNA-seq dataset generated for HeLa cells (Shen et al., 2014; Xue et al., 2013). This analysis indicated that upon PTBP1 knockdown, there were 289 significantly altered splicing events (FDR < 0.05, Figure 2A and Table S2), variously including cassette exons (CE), mutually exclusive exons (MXE), alternative 5’ splice sites (A5SS), alternative 3’ splice sites (A3SS), and intron retention (IR). Among the 215 altered CE events, PTBP1 apparently repressed 135 CEs and promoted the splicing of 80 CEs (Figure 2A). We next chose 30 PTBP1-repressed or -enhanced CEs for validation using semiquantitative reverse-transcriptase polymerase chain reaction (RT-PCR) with control or PTBP1 shRNA treated HeLa cells. We found that 90% of the examined splicing events showed consistent changes as detected by RNA-seq (Figures 2B, 2C, S2A, and S2B), suggesting the accuracy of the identified splicing events.

**Figure 2.**
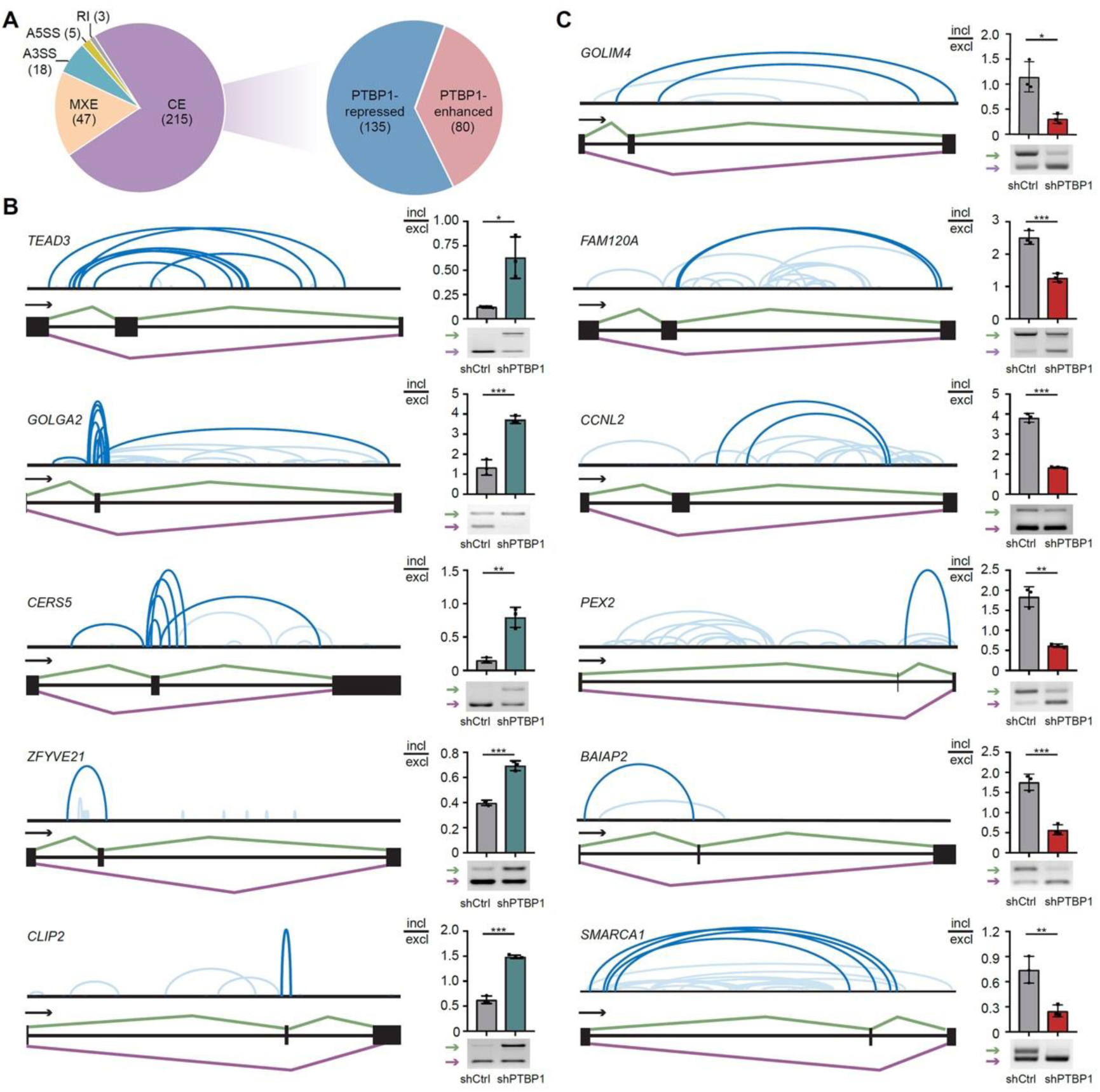
PTBP1-mediated RNA-RNA interactions regulate cassette exons splicing. (A) Pie charts showing PTBP1-regulated splicing events. CE, cassette exon. MXE, mutually exclusive exon. A5SS, alternative 5’ splice site. A3SS, alternative 3’ splice site. RI, intron retention. (B) Examples of PTBP1-mediated RNA-RNA interactions in repressing cassette exon splicing. The snapshots on the left exhibit CRIC-seq loops associated with splice sites (top, dark blue arc lines) and the PTBP1-regulated exon with its flanking intronic and exonic regions (exon, black box; intron, black line). Light blue arc lines denoted other chimeric reads around PTBP1-regulated exons. Green arrows indicate the long isoforms, and purple arrows indicate the short isoforms. The changes upon PTBP1 knockdown validated by semiquantitative RT-PCR are shown in the bar graphs on the right. Ctrl, control. (C) Examples of PTBP1-mediated RNA-RNA interactions in activating cassette exon splicing. Data in (B) and (C) are mean ± s.d.; n=3 biological replicates, two-tailed, unpaired *t*-test. \**P* < 0.05, ^∗∗^*P* < 0.01, ^∗∗∗^*P* < 0.001.

Previous studies have shown that PTBP1 regulates alternative splicing in a position-dependent manner (Llorian et al., 2010; Van Nostrand et al., 2020; Xue et al., 2009), but the underlying mechanism(s) remain unclear. We examined the PTBP1-mediated RNA looping patterns around the proximal and distal splice sites of the 215 identified CE splicing events. Specifically, loops with two arms mapped to the 500 nt intronic and the 50 nt exonic regions around exon-intron junction sites were retained for pattern searching. We ultimately found two general modes of RNA loops that modulate the splicing of CEs. Among the 135 PTBP1-repressed CEs, 119 of them contained PTBP1-mediated loops, and 65.5% of these were formed between the two surrounding introns (Figures 2B and S2A). In contrast, among the 69 PTBP1-activated CEs that contained PTBP1-mediated RNA loops, 94.2% of the RNA loops were enriched in introns (Figures 2C and S2B). These results suggest a position-dependent mode of PTBP1-mediated RNA loops in regulated splicing events.

Next, we constructed a composite 3D RNA functional map by aligning RNA loops within a 2.5-kb window around the splice sites of PTBP1-repressed and PTBP1-enhanced CEs on a scaled pre-mRNA model (Figures 3A). As controls, we built an RNA map for 7,181 constitutive exons (Figure 3A, right panel). Again, we observed a pronounced enrichment of exon-spanning RNA loops formed between the flanking intronic regions for PTBP1-repressed CEs (Figure 3A, middle panel). This result indicates that the previously proposed “looping out of exon” model may generally be applicable for explaining PTBP1-repressed CEs. In addition, a long-speculated “looping out of branch point” model (Liu et al., 2002) also seems to be correct, as 52.1% of the PTBP1-repressed CEs contained RNA loops around the 3’ splice sites and branched A points.

**Figure 3.**
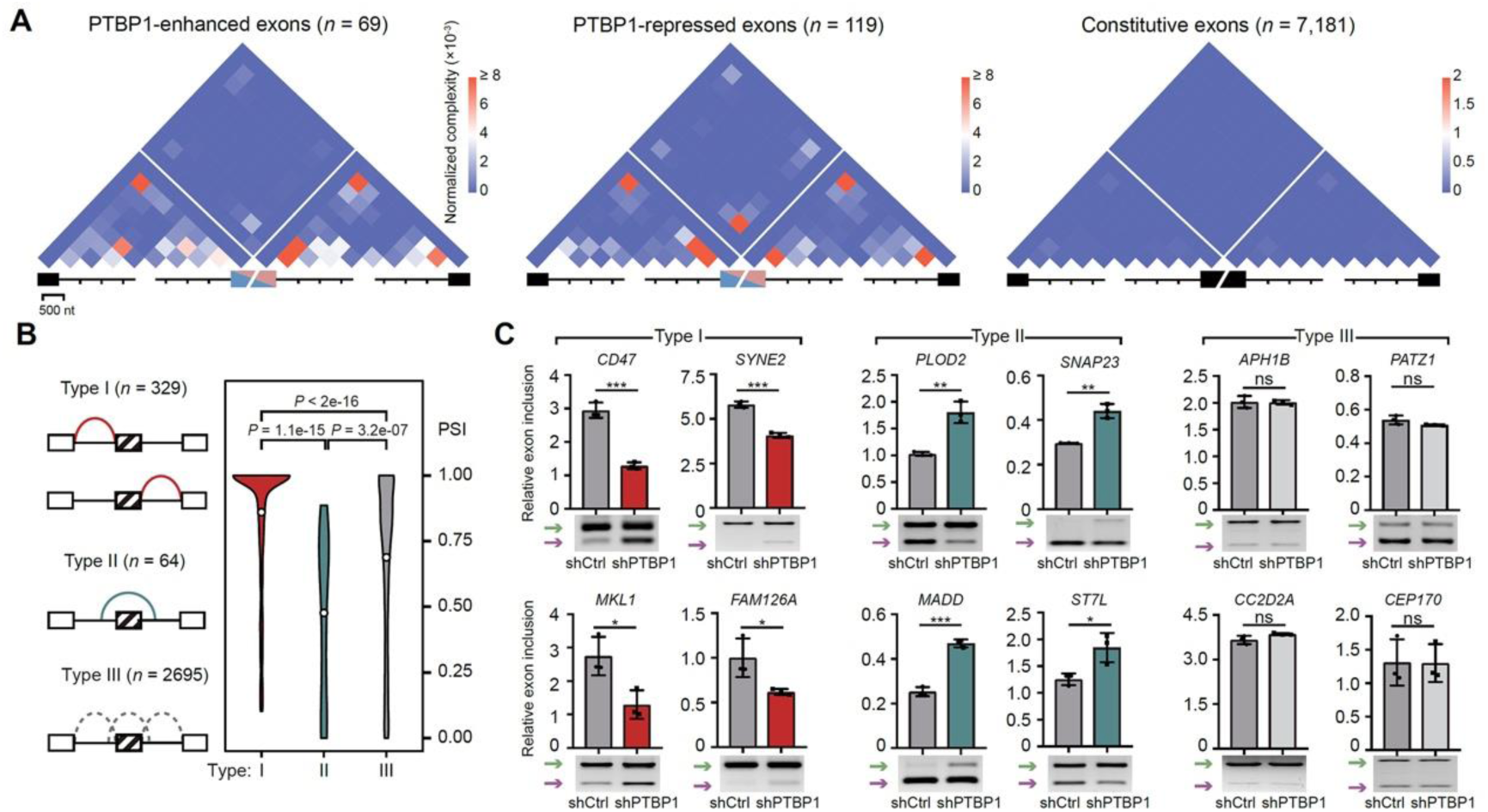
Positional rule of PTBP1-mediated RNA-RNA interactions in cassette exon splicing. (A) 3D RNA maps showing the positional rules for PTBP1-mediated cassette exon inclusion or exclusion. The RNA map for constitutive exons serves as a negative control and is shown on the right. (B) Violin plot showing the percent spliced in (PSI) values of PTBP1-activated (Type I), -repressed (Type II), and non-regulated (Type III) exons. Two-tailed unpaired *t*-test was used to calculate the *P* values. (C) RT-PCR validation of splicing events predicted based on the rules revealed by 3D RNA maps in (A). Green arrows indicate the long isoforms, and purple arrows indicate the short isoforms. Data are mean ± s.d.; n=3 biological replicates, two-tailed, unpaired *t*-test. ^ns^*P* >0.05, ^∗^*P* < 0.05, ^∗∗^*P* < 0.01, ^∗∗∗^*P* < 0.001.

For PTBP1-enhanced CEs, our 3D RNA functional map revealed predominant enrichment of RNA loops at flanking introns (Figures 3A, left panel). However, this mode did not mean that two flanking introns must simultaneously contain intronic RNA loops. Notably, 75.4% of the PTBP1-enhanced CEs showed the enrichment of RNA loops either in upstream or downstream introns. We thus speculated that the asymmetrical enrichment of RNA loops on one side of introns might be directly associated with splicing activation. Notably, PTBP1-mediated RNA loops were much less observed around 3’ splice sites and the branch A points in PTBP1-enhanced CEs than PTBP1-repressed CEs (Figure 3A). These results suggest that PTBP1 represses or enhances CE splicing through different RNA looping modes.

### RNA looping modes for predicting PTBP1 regulated splicing

In addition to the identified 215 PTBP1-regulated CEs (Figure 2A), we obtained 3,088 altered CEs (|ΔPSI| ≥ 0.1) with FDR values larger than 0.05 from the aforementioned RNA-seq dataset upon PTBP1 depletion. We next explored whether the revealed RNA looping modes could predict more PTBP1-regulated CEs from these CE inputs. We arbitrarily set the prediction criteria as follows: if the number of RNA loops (n≥4) at one side of introns were 4-fold more than those on the other side and also 4-fold more than the loops spanning across the CEs (see Type I, Figure 3B), we classified a given CE as a “PTBP1-enhanced event”. Conversely, if the number of RNA loops spanning across the CEs (n ≥ 4) was 4-fold more than the upstream and downstream intron-containing loops (see Type II, Figure 3B), a given CE was considered to be a “PTBP1-repressed event”. Using these criteria, we could predict 329 and 64 CEs that are most likely to be enhanced or repressed by PTBP1, respectively (Figure 3B and Table S3). The remaining 2,695 CEs (Type III) were kept as controls for comparison because no obvious PTBP1-mediated looping pattern was observed.

We next examined the percentage spliced in (PSI) differences for these three types of CEs to validate our prediction results (Figure 3B). In theory, the PTBP1-enhanced CEs in type I should have larger PSI values than PTBP1-repressed CEs in type II. Indeed, we found that the type I CEs had significantly larger PSI values than the type II CEs (*P*=1.1e-15, Figure 3B, right panel). As expected, the type III CEs (n=2,695) had the values in the middle (Figure 3B). These results indicate that PTBP1-mediated RNA loops can faithfully predict splicing outcomes. Next, we randomly chose 30 predicted CEs, including 22 CEs in type I and type II, for RT-PCR validation. As controls, 8 CEs in type III were also included for the analysis. Significantly, 90% of the tested events showed accordant changes as predicted (Figures 3C and S3), suggesting that PTBP1-mediated RNA looping modes are highly reliable in predicting new CEs regulated by PTBP1. The two CEs showing inconsistent changes with CRIC-seq prediction may be because the normalized loop numbers of these events are not enough for accurate predicting (Figure S3C, *ATG5* and *LRRC23*), which can be improved by increasing sequencing depth. Together, our CRIC-seq mapping and 3D RNA map analysis revealed a positional rule for PTBP1-mediated RNA loops in splicing regulation.

### PTBP1-mediated RNA loops facilitate asymmetrical intron removal to promote exon inclusion

Our CRIC-seq mapping and experimental validation confirmed that PTBP1 could repress splicing through “looping out of exon” or “looping out of branch point” mechanisms (Figure 3A). To explore how PTBP1-mediated intronic RNA loops enhance splicing, we determined to measure the removal rate for two surrounding introns, which tends to be asymmetrically removed with different rate and sequence (Gee et al., 2000; Kessler et al., 1993; Lang and Spritz, 1987; Tsai et al., 1980; Ule et al., 2006). We adapted an RT-PCR assay in shCtrl and shPTBP1 treated HeLa cells to quantify the splicing intermediates (SI) that contain one unspliced intron and the other exon spliced together (Figure 4A). After normalizing the products of 5’ SIs (F + Ra primer pair) or the 3’ SIs (Fa + R) to the pre-mRNA levels (Fa + Ra), we could quantify the two surrounding intron removal rates by dividing the 5’ SI or 3’ SI products to the products of pre-mRNA (Figure 4A).

**Figure 4.**
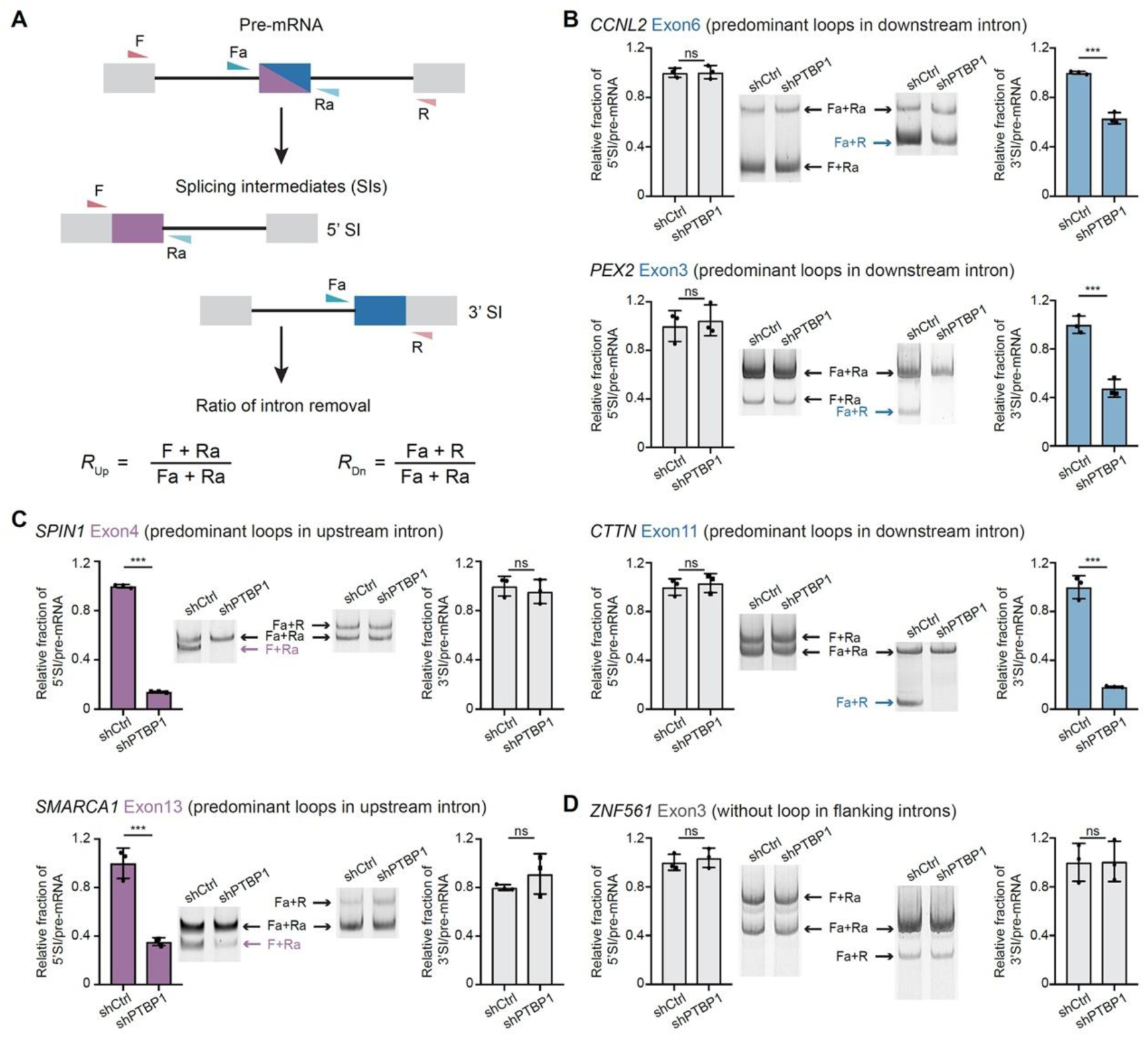
PTBP1-mediated asymmetrical RNA loops facilitate intron removal. (A) Schematic diagrams showing quantification of splicing intermediates by RT-PCR. Cassette exons are marked as blue and purple half rectangle triangles, constitutive exons are marked as gray boxes, introns are black lines, and primers are marked as colored triangles. Pre-mRNAs (up) are amplified by primers Fa and Ra; splicing intermediates (middle) are amplified by primers F and Ra (5’ splicing intermediates) or primers Fa and R (3’ splicing intermediates). (B) RT-PCR shows that PTBP1 intronic loops at the downstream intron of *CCNL2*, *PEX2*, and *CTTN* promote the intron removal rate. The blue or purple arrows indicate the splicing intermediates significantly changed upon PTBP1 knockdown, while the black arrows indicate unchanged splicing intermediates. The bar graphs quantify the relative amount of splicing intermediates normalized to pre-mRNAs. *R*_Up_ and *R*_Dn_ in shCtrl samples were normalized to 1. (C) RT-PCR shows that PTBP1 intronic loops at the upstream intron of *SPIN1* and *SMARCA1* promote the intron removal rate. (D) RT-PCR shows that the splicing intermediates of a PTBP1-unregulated *ZNF561* exon 3 were not changed upon PTBP1 knockdown. Data in (B), (C), and (D) are mean ± s.d.; n=3 biological replicates, two-tailed, unpaired *t*-test. ^ns^*P* >0.05, ^∗∗∗^*P* < 0.001.

Next, we randomly chose 13 PTBP1-regulated CEs for SI analysis. Among which, the SIs of five PTBP1-enhanced CEs (*CCNL2* exon 6, *PEX2* exon 3, *CTTN* exon 11, *SPIN1* exon 4, and *SMARCA1* exon 13) were analyzed by semi-quantitative PCR for subsequent agarose gel analysis. For PTBP1-mediated RNA loops located at the downstream introns, we observed significantly reduced 3’ SIs for all the three tested CEs (*CCNL2*, *PEX2*, *CTTN*) upon PTBP1 knockdown (Figure 4B). As an internal control, there were no differences between the 5’ SI levels in shCtrl and shPTBP1 treated cells (Figure 4B). Conversely, upon knocking down PTBP1, we observed significantly reduced 5’ SIs for PTBP1-mediated RNA loops located at the upstream introns. The 3’ SIs containing the downstream introns and unspliced exons were not changed (Figure 4C). As another control, we did not see differences in the SIs of a PTBP1 unregulated CE in *ZNF561* (Figure 4D). Moreover, we performed RT-qPCR to quantify the SIs for another 8 PTBP1-regulated CEs. Again, they all showed PTBP1-dependent and enhanced asymmetric intron removal (Figures S4A-D). These results suggest that PTBP1-mediated intronic RNA loops may enhance splicing by facilitating intron removal asymmetrically.

### PTBP1 dimerization-organized RNA loops are required for splicing regulation

Although the PTBP1 protein was characterized as a monomer in solution (Amir-Ahmady et al., 2005; Monie et al., 2005), two studies have demonstrated that it can also be present as a dimer in the nucleus (Gong et al., 2021; Xue et al., 2009). We next explored whether its dimerization contributes to PTBP1-mediated RNA loops detected in the CRIC-seq-datasets. We first performed a reciprocal immunoprecipitation assay with exogenously expressed Flag or HA-tagged PTBP1 in HeLa cells (Figure S5A). We found that the protein indeed could present as a dimer (Figure 5A). Moreover, the dimerization seems not dependent on RNA and DNA because combined treatment with RNase A and DNase I showed no influence on the pulled proteins (Figure 5A), indicating a physical association between two different PTBP1 molecules.

**Figure 5.**
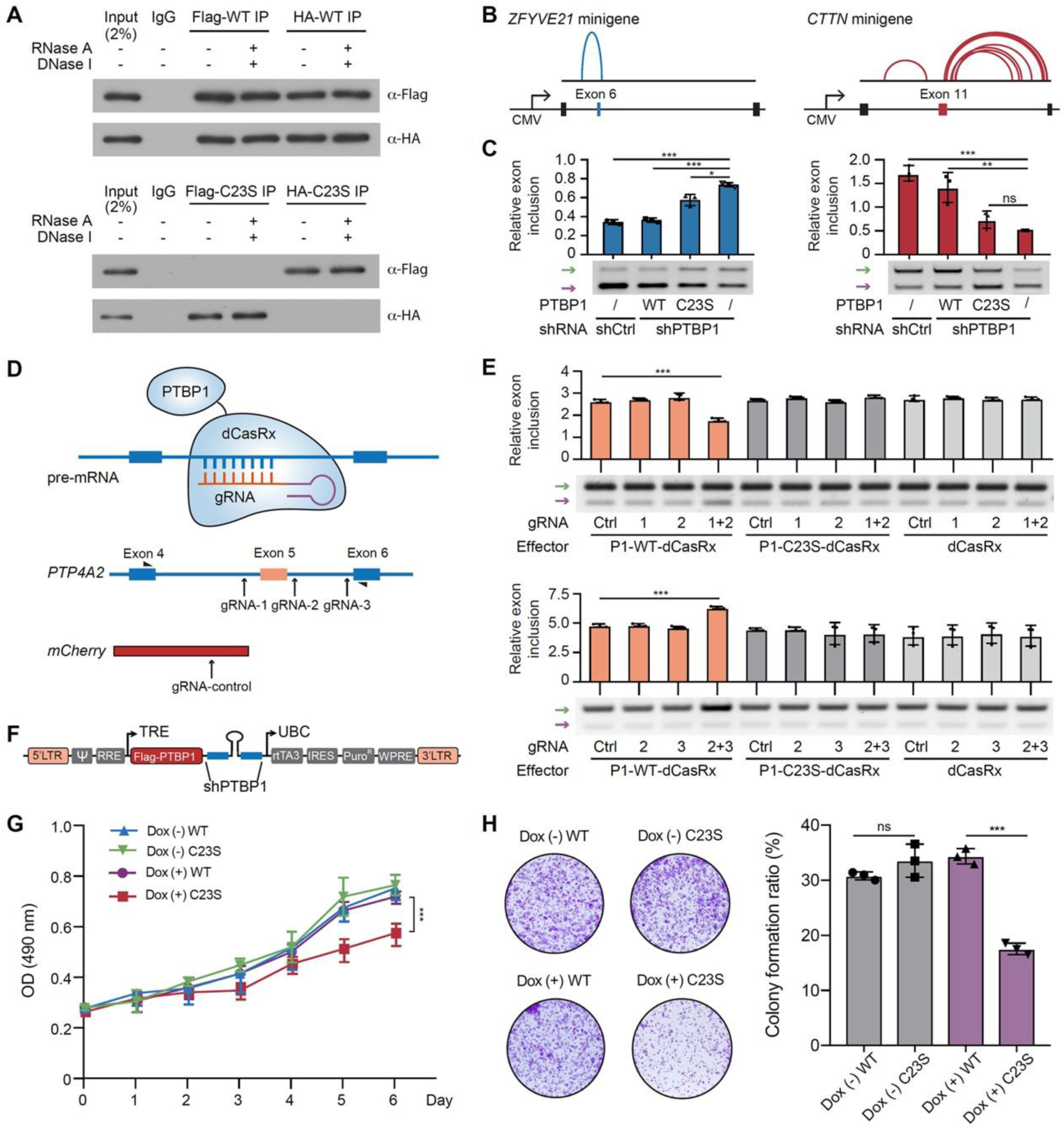
Disrupting PTBP1 dimerization-mediated RNA loops causes abnormal splicing isoforms and cell proliferation. (A) Co-IP showing that C23S mutant disrupts the dimerization between FLag- and HA-tagged PTBP1 proteins in transiently transfected HeLa cells. WT, PTBP1-WT protein; C23S, PTBP1-C23S protein. (B) Schematic diagram of the *ZFYVE21* and *CTTN* minigene reporters for validating PTBP1-dependent exon exclusion or inclusion, respectively. The splice site-associated RNA loops were shown in blue or red arc lines. (C) RT-PCR showing the splicing of *ZFYVE21* and *CTTN* minigene in response to PTBP1 knockdown with or without exogenously expressed PTBP1-WT or PTBP1-C23S protein. Green arrows indicate the long isoforms, and purple arrows indicate the short isoforms. (D) Diagram of gRNA-mediated dCasRx-PTBP1 or dCasRx-C23S fusion protein tethering assay to manipulate alternative splicing *in vivo*. Three gRNA targeting sites are marked by black arows as gRNA-1, gRNA-2, and gRNA-3. A gRNA targeting mCherry served as a negative control. (E) Semiquantitative RT-PCR shows gRNA-mediated and position-dependent exon inclusion or exclusion for endogenous *PTP4A2* pre-mRNA. Green arrows indicate the long isoforms, and purple arrows indicate the short isoforms. Ctrl, control; P1, PTBP1. (F) Schematic representation of pTRIPZ inducible lentiviral vector to simultaneously knock down endogenous PTBP1 and exogenously express shRNA-resistant PTBP1-WT or PTBP1-C23S. (G) Cell viability assay shows that the proliferation of PTBP1-C23S mutant HeLa cells is significantly decreased compared with wild-type (WT) controls. (H) The clonogenic assay shows that the C23S mutant significantly reduced the proliferation of HeLa cells. Data in (C), (E), (G), and (H) are mean ± s.d.; n=3 biological replicates, two-tailed, unpaired *t*-test. ^ns^*P* >0.05, ^∗^*P* < 0.05, ^∗∗^*P* < 0.01, ^∗∗∗^*P* < 0.001.

PTBP1 can form a homodimer in solution via a Cys23-mediated disulfide bond under non-reduced conditions (Monie et al., 2005). We found that the amino acid also contributed to the dimerization of PTBP1 within the cells because Flag- or HA-tagged C23S (cysteine to serine) mutant proteins failed to interact with each other (Figure 5A). In addition, our SDS-PAGE gel analysis showed that the purified wild-type protein could form a dimer, while the dimerization ability of the C23S mutants was largely disrupted (Figure S5B). Moreover, the gel shift assay revealed that the wild-type and the C23S mutant form of PTBP1 have comparable RNA binding activities (Figures S5C and S5D).

Next, we investigated whether PTBP1 dimerization could modulate CE splicing. We selected a PTBP1-repressed CE in *ZFYVE21* containing RNA loops across exon and a PTBP1-enhanced CE in *CTTN*, which has extensive intron spanning loops (Figure 1E), for detailed analysis by constructing two minigene reporters. After inserting the cassette exon (*ZFYVE21* exon 6 or *CTTN* exon 11) and its flanking introns and exons into the pcDNA3 backbone (Figure 5B), we transiently transfected these two reporters into shCtrl, or shPTBP1 treated HeLa cells (Figure 5C). For comparisons, we also co-transfected an shRNA-resistant wild-type PTBP1 or C23S mutant, both of which contained synonymous mutations at shRNA targeting sites.

Upon PTBP1 knockdown, the splicing of exon 6 in the *ZFYVE21* minigene was significantly increased compared to control shRNA-treated cells (Figures 5C and S5E, left panel, lane 1 vs. lane 4). However, expression of a wild-type form of PTBP1 in shPTBP1-treated HeLa cells could repress the exon 6 splicing to the level of shCtrl-treated samples (Figures 5C and S5E, left panel). In contrast, the C23S mutant proteins failed to suppress the exon 6 splicing (Figures 5C and S5E, left panel). These results indicate that PTBP1 dimerization can induce RNA looping across CEs to repress its splicing. For the *CTTN* minigene reporter, we found that the inclusion of the exon 11 was significantly reduced in response to PTBP1 knockdown (Figures 5C and S5E, right panel, lane 1 vs. lane 4). Moreover, the reduced usage of exon 11 could be rescued by the wild-type PTBP1 but not the C23S mutant (Figures 5C and S5E, right panel), highlighting that PTBP1 dimerization-induced RNA loops within introns can enhance CE splicing. Together, these results suggest that PTBP1 dimerization is required for long-range loop formation and is directly involved in splicing regulation.

### PTBP1 dimerization drives RNA loop formation to modulate splicing

To see whether PTBP1 dimerization-mediated RNA looping can be artificially engineered to regulate CE splicing, we adopted an RNA-targeting CRISPR-dCasRx strategy (Du et al., 2020; Konermann et al., 2018). In this tethering assay, wild-type or the C23S mutant PTBP1 was fused to the N-terminus of dCasRx and then was artificially tethered to specific intronic regions by guide RNAs (gRNAs) (Figure 5D). Using this system, we could tether the PTBP1-dCasRx to one side or both sides of the intron using two different pairs of gRNAs to induce RNA loop formation. As controls, the C23S-dCasRx lacking dimerization abilities was also included for comparisons. In addition, a guide RNA targeting to mCherry was included as another negative control (gRNA-control, gRNA-Ctrl). Moreover, to exclude the off-target effects of the CRISPR-dCasRx system, the dCasRx protein without PTBP1 fusion was also introduced as an additional negative control (Figure 5D).

We chose a PTBP1 non-target gene *PTP4A2* (exon 5) for the tethering analysis. Three gRNAs were designed: one to target the upstream 3’ splice site (gRNA1), one for the downstream 5’ SS (gRNA2), and one for the downstream 3’ SS (gRNA3) (Figure 5D). The gRNAs were individually expressed or combined (gRNA1+2 and gRNA2+3) for tethering dCasRx protein or PTBP1-dCasRx fused protein to the targeted regions. We found that the inclusion of exon 5 was reduced by 35% after combined transfection of gRNA1+2 and PTBP1-dCasRx in HeLa cells (Figure 5E, up panel, lane 1 vs. lane 4; Figure S5F, left panel). Notably, the effects were not due to ectopically expressed dCasRx because the control gRNA (to mCherry) group had no influence on exon 5 splicing (Figure 5E, up panel; see lanes 1 and 9-12). In contrast to the wild-type PTBP1, the C23S-dCasRx mutant proteins failed to suppress the splicing of exon 5 (Figure 5E). These results demonstrate that engineered PTBP1 RNA loops across CEs can repress its splicing.

We next tested whether engineered PTBP1 RNA loops in introns can enhance CE splicing. By tethering PTBP1-dCasRx to the downstream intron 5 of *PTP4A2* with gRNA2 and gRNA3 simultaneously, we observed an approximately 30% increase in the inclusion of *PTP4A2* exon 5 (Figure 5E, bottom panel, lane 1 vs. lane 4; Figure S5F, right panel). In contrast, individually tethering PTBP1-dCasRx with either gRNA2 or gRNA3 did not change the distribution of splice isoforms. Nor were any changes in the exon 5 inclusion detected upon the expression of C23S-dCasRx or dCasRx combined with a control gRNA to mCherry (Figure 5E, bottom panel; Figure S5F, right panel). These results show that PTBP1 dimerization-mediated RNA looping is required for both splicing activation and repression in a position-dependent manner.

### Splicing QTLs disrupt PTBP1-mediated RNA loops to drive tumorigenesis

PTBP1 tends to be upregulated in various tumors (David et al., 2010; He et al., 2014; He et al., 2007). We next investigated whether its dimerization has any influence on tumor cell proliferation. We here adopted a doxycycline (Dox)-inducible replacement strategy by knocking down PTBP1 with a previously validated shRNA (Xue et al., 2009), and by simultaneously expressing an shRNA-resistant form of PTBP1 or C23S mutant (synonymous mutation in shRNA targeting site, Figures 5F and S5G). With this system, in the presence of doxycycline, either shRNA-resistant PTBP1 or C23S mutant gradually replaces the endogenous PTBP1 population. We found that the C23S mutant significantly reduced HeLa cell proliferation (Figures 5G and 5H), indicating that PTBP1 dimerization is required for tumor cell growth.

To further examine how PTBP1 dimerization regulates tumorigenesis, we analyzed disease-associated genetic variants that might disrupt PTBP1-mediated RNA loops in pre-mRNA. Among the vast amounts of GWAS loci, we focused on splicing quantitative trait loci (sQTLs) known from the literature to be directly associated with alternative splicing. We analyzed the 819,254 sQTLs collected by the CancerSplicingQTL database (Tian et al., 2019), of which, 146,426 sQTLs were directly overlapped with the 271,286 clusters of PTBP1-mediated loops (Figure 6A and Table S4). Among the 581 PTBP1-bound and regulated CE splicing events defined by RNA-seq and CRIC-seq, 121 of them contained at least one sQTL in PTBP1-mediated RNA loops, suggesting that these sQTLs may affect splicing by altering these loops.

**Figure 6.**
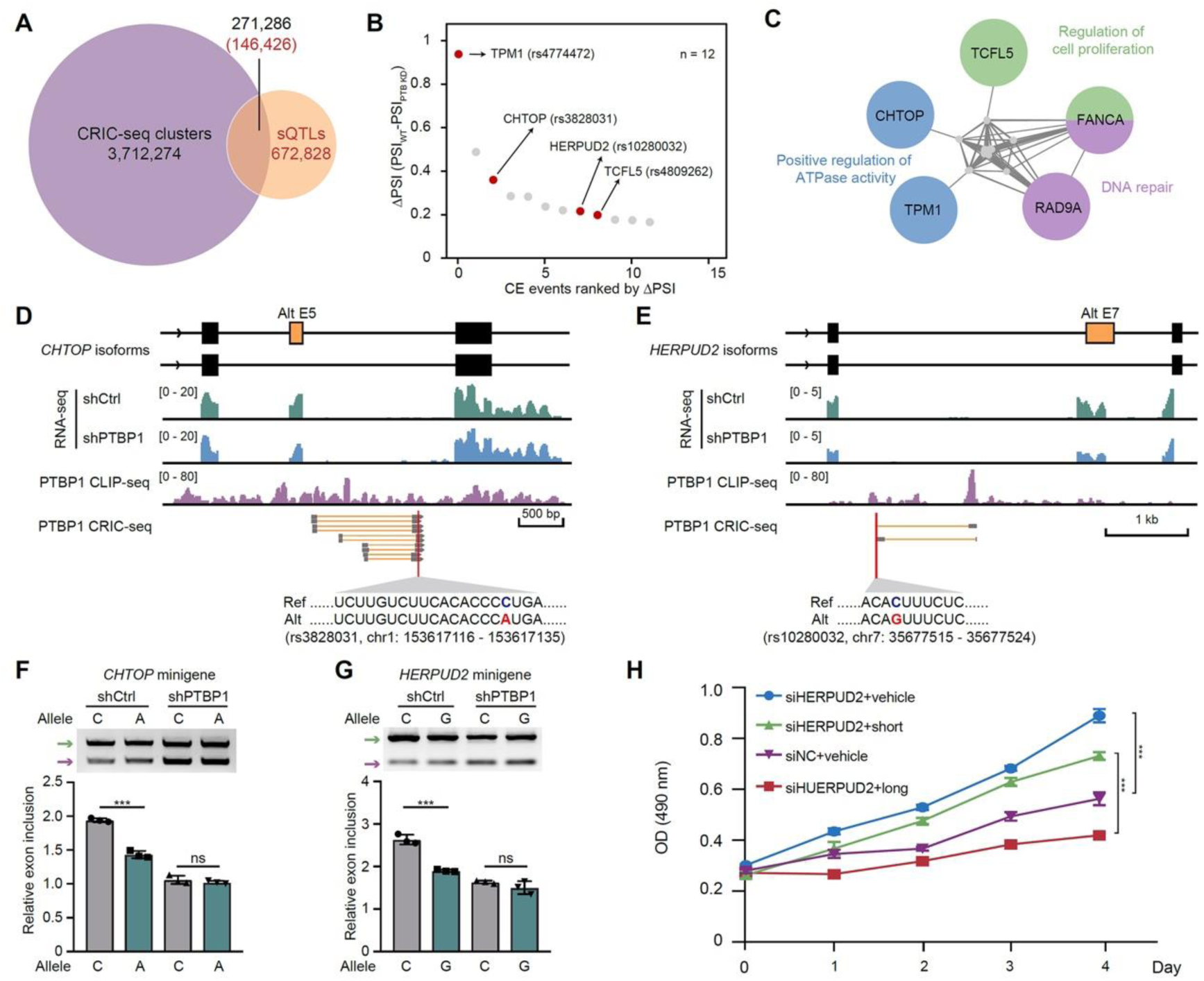
Splicing quantitative trait loci (sQTL) disrupt PTBP1-mediated loops to induce tumorigenesis. (A) Venn diagram showing cancer-related sQTLs overlapped with PTBP1-mediated CRIC-seq clusters. (B) Scatter plot showing the 12 sQTLs that may influence cassette exon splicing. (C) Protein network analysis for the sQTLs affected 11 genes. Several genes associated with Gene Ontology terms were listed in different colors. (D, E) Snapshot of PTBP1 CLIP-seq and CRIC-seq signals in *CHTOP* and *HERPUD2* cassette exons and their flanking introns and exons. RNA-seq signals for shCtrl and shPTBP1-treated HeLa cells are shown in green and blue, respectively. CLIP-seq reads are shown in purple. CRIC-seq clusters containing sQTLs (red) are shown in gray boxes and magnified at the bottom. (F, G) The splicing of sQTLs containing *CHTOP* and *HERPUD2* reporters showed decreased exon usage in shCtrl-treated HeLa cells, and the mutants failed to respond to PTBP1 knockdown. Green arrows indicate the long isoforms, and purple arrows indicate the short isoforms. (H) Cell viability assay shows that the longer isoform of *HERPUD2* represses cell proliferation and the shorter isoform has opposite effects. Data in (F) and (H) are mean ± s.d.; n=3 biological replicates, two-tailed, unpaired *t*-test. ^ns^*P* >0.05, ^∗∗∗^*P* < 0.001.

To examine how these sQTLs affect the binding affinity of PTBP1, we first performed an *in vitro* systematic evolution of ligands by exponential enrichment (SELEX) assay to quantify the binding affinity of PTBP1 on various RNA sequences. From the initial pool of 1,048,576 random RNA sequences, we did three rounds of binding and RNA pull-down with a low concentration of his-tagged PTBP1 protein by following a previously published protocol (Lambert et al., 2014). The enriched RNAs were purified and converted into a library for sequencing (Figure S6A). Using the enriched RNA sequences, we deduced a classical PTBP1 binding motif CCUCUCC (Figure S6B), indicating that our SELEX results are reliable. Next, we rank the enriched RNA sequences based on their sequencing read number, which can relatively represent the binding preference of PTBP1 to the sequence. By aligning the sQTL-containing sequences to the ranked sequences revealed by SELEX, we quantified the relative influence of each sQTL on PTBP1 binding affinity (Figure S6C).

As PTBP1 preferentially binds CU-rich elements (Figure S6B), we focused on the sQTL transversions that changed A/G to C/U or vice versa. Overlapping analysis with the PTBP1-regulated CEs enabled us to identify 12 high-confidence splicing events that might be influenced by cancer-related sQTL transversions (Figures 6B and S6C). We randomly selected four CEs, including the exon 5 of *CHTOP* gene, exon 7 of *HERPUD2*, exon 6 of *TPM1*, and exon 4 of *TCFL5*, for validation in shPTBP1-treated HeLa cells by RT-PCR (Figure 6B, red dots). We found that these four CEs showed PTBP1-depdent splicing activation compared with shCtrl-treated samples (Figure S6D). For the 12 sQTLs-associated 11 genes, we analyzed their protein-protein interaction networks. We found that five affected genes, *CHTOP*, *TPM1*, *TCFL5*, *RAD9A*, and *FANCA*, showed some shared interaction partners and were involved in several biological processes, including DNA repair, regulation of cell proliferation, and ATPase activity (Figure 6C).

To dissect the mechanism by which these sQTL transversions affect splicing, we constructed four minigene reporters for *CHTOP*, *TCFL5*, *TPM1* and *HERPUD2* genes by cloning their CEs and flanking introns as well as constitutive exons into the pcDNA3 backbone (Figures 6D, 6E, S6E, and S6F). According to the sQTLs, we also made relevant mutations on reporters for comparisons. Notably, the surrounding introns of these four CEs contained PTBP1-mediated RNA loops. Based on our *in vitro* SELEX data, the sQTLs (C-to-A or C-to-G) in the surrounding introns of these CEs were predicted to reduce the binding affinity of PTBP1 and cause decreased exon inclusion levels. Next, we transfected the wild-type or sQTL-containing reporters into shCtrl or shPTBP1-treated HeLa cells. Similarly, PTBP1 also enhanced the splicing of these four exons in minigene reporters (Figures 6F, 6G, S6G, and S6H). However, mutation of the reference into sQTL-associated nucleotides reduced the CEs splicing. Importantly, the sQTLs introduced into minigene reporters almost disrupted the regulation of PTBP1 on these CEs (Figures 6F, 6G, S6G, and S6H).

Next, we determined to investigate which sQTL altered CE splicing event directly contributes to PTBP1 dimerization-mediated proliferation advantages in HeLa cells (Figures 5G and 5H). Among the tested four genes *CHTOP*, *TCFL5*, *TPM1* and *HERPUD2*, only the expression level of *HERPUD2* is correlated with the survival probability of cervical cancer patients by checking The Human Protein Atlas database. We therefore focused on *HERPUD2* for detailed phenotypic analysis using cervical cancer cell line HeLa. We found that knocking down *HERPUD2* enhanced HeLa cell proliferation by 2 fold, and the phenotypic alteration could only be reversed by expression of the exon 7-containing *HERPUD2* isoform rather than the shorter isoform without exon 7 (Figure 6H). This result might explain why the sQTL (C-to-G transversion) induced exon 7 skipping in the *HERPUD2* could promote tumorigenesis (Figure 6H). Taken together, our results indicate that cancer-related sQTLs could disrupt PTBP1-mediated RNA loops to induce cancer and other diseases.

## DISCUSSION

Many RBPs have been demonstrated to be co-transcriptionally interacted with pre-mRNAs via specific motifs to regulate splicing outcomes in a position-dependent manner (Fu and Ares, 2014). However, how the positional effects are achieved remains unclear. In this study, we sought to tackle this problem by developing a CRIC-seq method to profile single RBP-mediated RNA-RNA spatial interactions globally. Applying CRIC-seq to map PTBP1-mediated long-range RNA loops in HeLa cells and the combined analysis of PTBP1-regulated splicing events enabled us to generate a 3D RNA map to investigate the mechanisms of positional effects from a new angle of RNA spatial conformation. We demonstrated that PTBP1-mediated “loop out of the exon” model is generally present and appears to be a major way to repress the splicing of CEs (Figure 7A). 3D RNA map also revealed a general principle for PTBP1-mediated splicing activation, in which the long-range RNA loops were asymmetrically localized in one surrounding intron to facilitate the intron removal rate (Figure 7B). Furthermore, we demonstrated that some cancer-related sQTLs might reduce the binding affinity of PTBP1 to the intronic region of pre-mRNAs and disrupt RNA loops mediated by the protein (Figure 7B), leading to significantly reduced exon inclusion of *CHTOP*, *TPM1*, *TCFL5*, and *HERPUD2*. Notably, the sQTL in *HERPUD*2 promotes the generation of a short splicing isoform that confers a growth advantage to cancer cells. Our results demonstrated the power of CRIC-seq in detecting a specific protein-mediated RNA loop and highlighted the crucial role of RNA conformation in splicing regulation and tumorigenesis.

**Figure 7.**
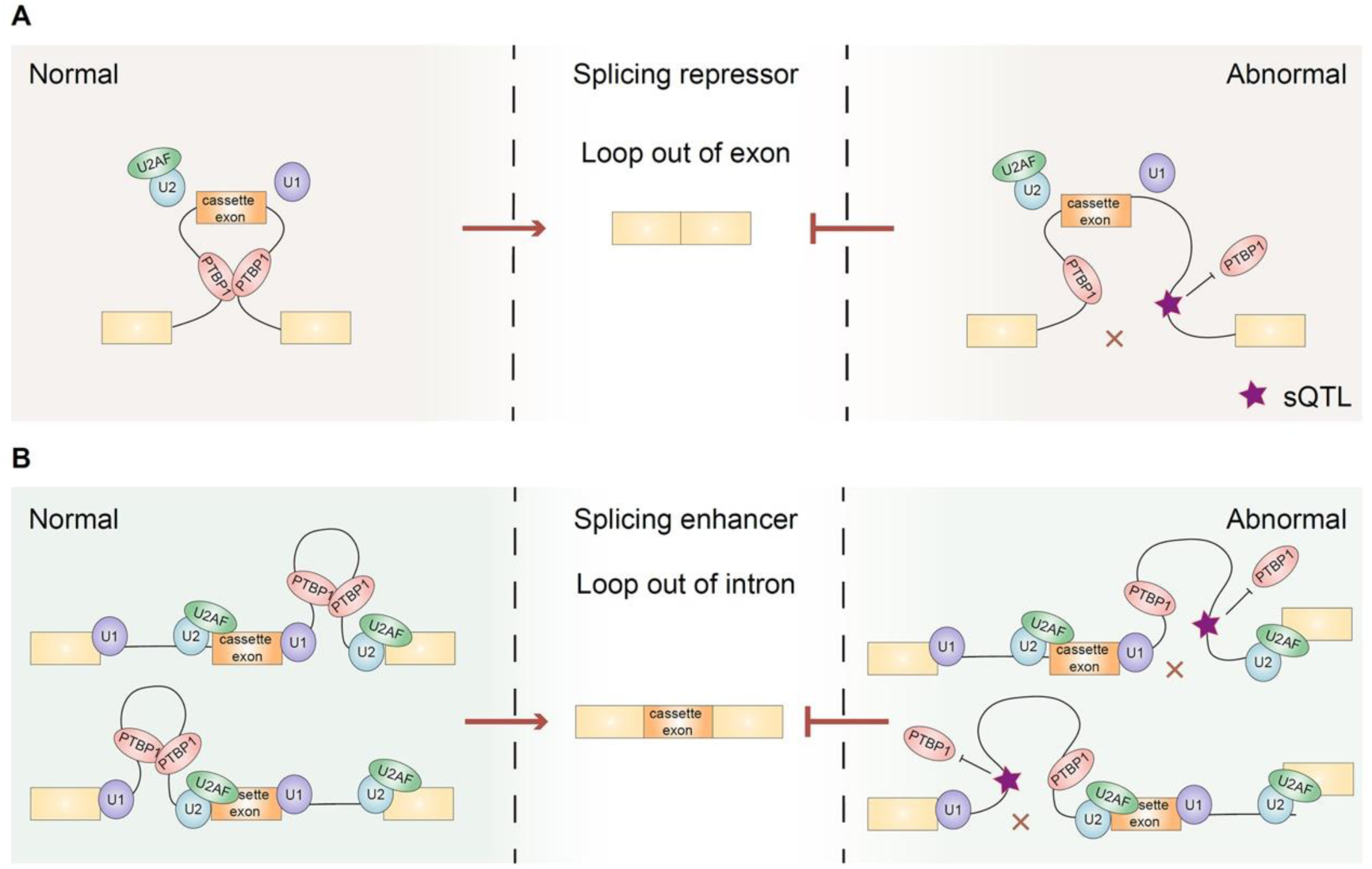
PTBP1-mediated long-range RNA looping in position-dependent alternative splicing and diseases. (A) PTBP1 binds to flanking introns and forms long-range RNA loops across the cassette exon to repress splicing. This model is consistent with a long-standing “loop out of exon” model. (B) PTBP1-mediated RNA loops, either in upstream or downstream introns, can promote the splicing of the cassette exon. This “loop out of intron” model can facilitate intron removal on one side asymmetrically. In these two kinds of models, cancer-related splicing QTLs located at the loop regions may lead to abnormal splicing by reducing the binding affinity of RNAs to PTBP1.

Mapping of RBP occupancy by CLIP-seq, eCLIP, and LACE-seq has enabled the generation of functional RNA maps for hundreds of splicing factors (Konig et al., 2010; Llorian et al., 2010; Ule et al., 2006; Van Nostrand et al., 2020; Wang et al., 2010; Xue et al., 2009). Although these 1D RNA binding maps established the correlation of binding to the functional consequence of the protein ablation, the underlying mechanisms remain poorly understood. As RNA molecules tend to fold into intricate structures with the help of various RBPs, therefore understanding regulated splicing from the RNA structure angle is needed. By mapping PTBP1-mediated RNA loops with CRIC-seq, we generated a first 3D RNA functional map to explain the positional effects of PTBP1 on regulated splicing (Figure 3A). We demonstrated that PTBP1 dimerization can drive long-range RNA loop formation in introns, further bringing the distal splice sites into proximity to promote splicing occurrence at the proximal sites. In such a scenario, we speculate that the PTBP1-mediated splicing outcome may strictly depend on where the RNA loops are formed. Suppose the loops formed between the upstream and downstream introns, the upstream 5’SS and downstream 3’SS were placed in proximity to promote exon skipping. In contrast, if the loops were formed either upstream or downstream of the intron, only one side of the splice sites in proximity was quickly removed and thus promoted splicing of the CEs.

Many RBPs can present as dimers or oligomers in the nucleus, such as PTBP1, hnRNPA1, and hnRNPK (Backe et al., 2005; Ding et al., 1999; Xue et al., 2009). We showed that the homotypic interaction between PTBP1 is required for maintaining over 10 million RNA loops, and many of them were involved in splicing regulation. Notably, PTBP1 could interact with other RBPs that can directly bind pre-mRNAs via specific motifs (Coelho et al., 2015; Gooding et al., 2013; Hahm et al., 1998; Huttelmaier et al., 2001). Therefore, some CRIC-seq detected RNA loops may be indirectly mediated by PTBP1-associated proteins. This situation partially explained why in the type I and type II loops we failed to see PTBP1 binding evidence on both sides or one side of the chimeric RNA arms (Figure S1). Of note, we used formaldehyde as a cross-linking reagent in our CRIC-seq approach to fix protein-mediated RNA-RNA interactions, which enabled us to detect both PTBP1 directly and indirectly interacting RNAs. It is conceptually different from CLIP-seq and various variant methodologies, which only detect direct binding sites of an RBP and lack the conformation information between these interacting sites.

RNA-RNA interactions tend to be mediated or stabilized by specific RBPs and play critical roles in regulating pre-mRNA splicing, translation, and RNA stability. Mapping of defined RBP-mediated inter-or intra-molecular RNA-RNA interactions appears to be a prerequisite for understanding their functional mechanisms in gene regulation. Several methods have been developed for such purpose, including two prime examples of CLASH and hiCLIP (Kudla et al., 2011; Sugimoto et al., 2015), which need the ectopic expression of a tagged protein in transfected cells for affinity purification and the subsequent proximity ligation *in vitro*. These methods have been successfully applied to identify the *in vivo* targets of snoRNAs, miRNAs, and piRNAs (Helwak et al., 2013; Kudla et al., 2011; Shen et al., 2018). However, the ectopic expression of a tagged protein may disrupt the native RNA-RNA interaction networks, and the proximity ligation in a diluted solution may lead to a higher false-positive rate. These limitations restrict their applications in studying the role of RNA spatial conformations in regulated splicing. We recently developed an RNA in situ conformation sequencing technology, RIC-seq, for profiling all the RBP-mediated RNA-RNA spatial interactions in situ (Cai et al., 2020). In this study, we further devised a captured RIC-seq method to identify a single RBP-mediated RNA-RNA interactions in native conditions. We speculate that CRIC-seq would be beneficial in understanding the functional mechanisms of other RBPs in various biological contexts.

GWAS studies have identified vast genetic variants localized in noncoding regions that may directly influence alternative splicing (Tam et al., 2019). In this study, we focused on sQTLs associated with various cancers to examine whether they could disrupt PTBP1-mediated long-range RNA loops and lead to aberrant pre-mRNA processing (Tian et al., 2019). By minigene-based analysis, we showed that these sQTLs indeed influence exon inclusion or exclusion by affecting loop formation via altering the PTBP1 binding affinity. It bears emphasis that we merely examined the splicing events significantly changed upon PTBP1 depletion, and the regulated events must contain multiple CRIC-seq reads. These strict criteria can plausibly explain that we only detected tens of sQTLs which can disrupt PTBP1-mediated RNA loops. It will be engaging in the future to evaluate all the cancer-associated sQTLs that potentially influence the PTBP1-mediated RNA loops and further quantify their effects using cancer genome sequencing data and the corresponding RNA-seq data.

Our studies also suffered from some limitations. For example, our analysis neglected approximately 1.8 million CRIC-seq identified intermolecular RNA-RNA interactions because they may not be directly involved in splicing regulation. It will be interesting in the future to see how this fraction of RNA-RNA interactions contributes to gene regulation. In addition, the faithful splicing of a CE tends to be regulated by multiple RBPs, which may be associated with each other or have antagonizing roles. Constructing combinatorial 3D RNA maps by integrating CRIC-seq data for multiple RBPs would likely help deepen understanding the complexity and specificity of regulated splicing. Moreover, CRIC-seq method is unable to discriminate RBP-mediated direct and indirect RNA loops, and integration analysis with CLIP-seq data in the same cell types is necessary to address the problem.

## ACKNOWLEDGMENTS

This work was supported by the National Natural Science Foundation of China (32130064, 32025008, 91940306, and 81921003), the National Key Research and Development Program of China (2017YFA0504400), and the Strategic Priority Program of CAS (XDB37000000), and the K.C. Wong Education Foundation (GJTD-2020-06) to Y.C.X., by the National Natural Science Foundation of China (31900465 and 32070620), and the National Key Research and Development Program of China (2021YFE0114900) to C.C.C.

## AUTHOR CONTRIBUTIONS

Y.C.X. conceived and supervised the project; R.Y. cultured cells, developed CRIC-seq method, create PTBP1 CRIC-seq libraries, and performed most of the experiments with the help from C.Y and S.H.X; R.B.S. performed *in vitro* SELEX assay; N.J.H. performed the bioinformatics analysis with the help from C.C.C.; Y.C.X. wrote the manuscript with the help of R. Y. and N.J.H.

## DATA AVAILABILITY

CRIC-seq sequencing data generated in this study are available in the GEO under accession number GSE210583.

## CODE AVAILABILITY

Codes used for CRIC-seq data analysis in this paper can be found at https://github.com/HuNaijing/CRIC-seq.

## SUPPLEMENTARY TABLES

Supplementary Table 1. Summary of the CRIC-seq libraries generated in this study.

Supplementary Table 2. List of PTBP1 regulated splicing events.

Supplementary Table 3. List of predicted splicing events regulated by PTBP1.

Supplementary Table 4. List of the overlapped sQTLs in PTBP1 CRIC-seq clusters.

Supplementary Table 5. List of primers and siRNAs used in this study.

## METHODS

### Cell culture

HeLa (ATCC, CCL-2) cells were cultured in a DMEM medium (Thermo Fisher, C11965500BT) containing 10% fetal bovine serum (PAN-Biotech, P30-3031) and 100 U/ml penicillin/streptomycin (Life Technologies, 15140) at 37 °C in 5% CO2 incubator. PCR routinely checked the HeLa cells to ensure they were free of mycoplasma contamination.

### Construction of CRIC-seq libraries

CRIC-seq library construction mainly includes crosslinking, permeabilization, MNase digestion, pCp-biotin labeling, proximity ligation, cell lysis, immunoprecipitation, RNA purification, DNase I treatment, RNA fragmentation, C-biotin selection, and strand-specific library preparation. Crosslinking, permeabilization, MNase digestion, pCp-biotin labeling and proximity ligation were performed as previously described (Cai et al., 2020), followed by the procedures:

#### Beads preparation and antibody coupling

During proximity ligation, protein A/G magnetic beads for immunoprecipitation were first washed three times with 0.1 M Na-phosphate buffer, pH 8.1 and then resuspended with 0.1 M Na-phosphate, pH 8.1 buffer containing the specific antibody in a fresh 1.5-ml Eppendorf tube. The tube was incubated at room temperature for 1 h by rotating at 20 rpm and then washed three times with wash buffer (1 × PBS with no Mg^2+^ and no Ca^2+^, 0.1% SDS, 0.5% deoxycholate, 0.5% NP-40). Anti-PTBP1 antibody (BB7) (Chou et al., 2000), anti-hnRNPA1 antibody (Santa Cruz, sc-32301), and anti-IgG antibody (Santa Cruz, sc-2025) were used.

#### Cell lysis and immunoprecipitation

After proximity ligation, cells were washed three times with 1 × PNK buffer (50 mM Tris-HCl pH 7.4, 10 mM MgCl_2_, 0.2% NP-40).The cell pellets were resuspended in wash buffer and then sonicated with BRANSON SLPe at 40% amplitude for 10 sec on/off for 3 cycles. After further incubation on ice for 15 min, the cell lysates were centrifuged at 13,000 rpm for 15 min at 4°C, and the supernatant was transferred to the Eppendorf tube that contains antibody-coupled beads. After gently mixing, the mixture was incubated at 4°C for 4 h by rotating at 20 rpm.

#### RNA purification

To elute RNA from the beads, 200 μl of proteinase K buffer (10 mM Tris-HCl pH 7.5, 10 mM EDTA, 0.5% SDS) and 50 μl of proteinase K (Takara, 9034) were added. The tubes were incubated at 37 °C for 60 min and 56 °C for 15 min, and the RNA was then extracted with 750 μl of TRIzol LS (Thermo Fisher, 10296028) and 220 μl of chloroform (Sinopharm, 10006818) following the manufacturer’s instructions. After centrifuging at 13,000 rpm for 15 min at 4 °C, the aqueous phase was transferred to a new 1.5 ml Eppendorf tube, added with 500 μl of 2-Propanol (Sigma, I9516-500ML) and 1 μl of glycoblue (15 mg/ml, Thermo Fisher, AM9515) and precipitated overnight at -20 °C. RNA was pelleted at 13,000 rpm for 20 min at 4 °C, washed twice with 75% ethanol, and dissolved in 15 μl of nuclease-free water.

#### DNase I treatment

To remove the potential joining of DNA to RNA mediated by T4 RNA ligase, 15 μl of RNA was treated with 10 μl of 10 × RQ1 DNase I buffer, 3 μl of RiboLock RNase Inhibitor (Thermo Fisher, EO0381) and 5 μl of DNase I (Promega, M6101) at 37 °C for 20 min. RNA was purified with ZYMO RESEARCH RNA Clean and Concentrator-5 (Zymo Research, R1016).

#### RNA fragmentation

To fragment RNA, 16 μl of total RNA was added with 4 μl of 5 × first-strand buffer (250 mM Tris-HCl pH 8.3, 375 mM KCl, 15 mM MgCl_2_), incubated at 94 °C for 5 min and immediately put on ice.

#### C-biotin selection

To enrich pCp-biotin-labeled chimeric RNAs, 20 μl of MyOne Streptavidin C1 beads (Thermo Fisher, 65002) were first washed twice with solution A (0.1M NaOH, 0.05M NaCl) and once with solution B (0.1M NaCl) and then blocked with yeast RNA (Roche, 10109223001). The blocked C1 beads, 30 μl of nuclease-free water, and 50 μl of 2 × TWB buffer (10 mM Tris-HCl pH7.5, 1 mM EDTA, 2 M NaCl, 0.02% Tween 20) were applied to the fragmented RNA. The mixture was incubated by rotating at RT for 30 min, after which the beads were washed four times with 1 × TWB buffer. To elute RNA from C1 beads, the beads were resuspended in 100 μl of PK buffer (100 mM NaCl, 10 mM Tris-HCl pH 7.0, 1 mM EDTA, 0.5% SDS) and incubated in Thermomixer at 95 °C for 10 min at 1000 rpm. Then the sample was placed on a DynaMag-2 magnet stand (Invitrogen, 12321D) for 1 min, and the aqueous phase was transferred to a new Eppendorf tube. Repeat the elution process once, and rinse the beads with 100 μl PK buffer. Finally, RNA was extracted from the 300 μl eluted sample with phenol chloroform 5:1 solution (Amresco, E277-400ML) and precipitated overnight at a final concentration of 300 mM NaCl, with 3 volumes of ethanol and 1 μl of glycoblue at - 20 °C. RNA was pelleted at 13,000 rpm, washed twice with 75% ethanol and dissolved in 10 μl of nuclease-free water.

#### Strand-specific library preparation

Strand-specific library construction for paired-end high throughput sequencing was performed as previously described (Cai et al., 2020). Briefly, first-strand cDNA was synthesized with random hexamer primer, and second-strand cDNA was synthesized by introducing dUTP in the dATP, dTTP, dCTP, and dGTP mix to achieve strand-specificity. Then the cDNA was performed with end-repair, dA-tailing, and adaptor ligation steps, followed by USER enzyme (NEB, M5505S) treatment and PCR amplification.

### Validation of PTBP1-regulated splicing events

Lentiviral shRNA against non-target and human *PTBP1* were packaged as previously described (Xue et al., 2013). HeLa cells were infected with lentiviral particle solution containing 8 μg/ml polybrene (Sigma, TR-1003-G) for 24 h followed by selection with hygromycin B (Gibco, 10687010) at a final concentration of 50 μg/ml for 48 h. Total RNA of selected HeLa cells was extracted with TRIzol Reagent (Invitrogen, 15596026) according to the manufacturer’s instructions. For RT-PCR, RNA was treated with RQ1 RNase-free DNase (Promega, M6101) and reverse transcribed with random hexamer primers using MMLV reverse transcriptase (Promega, M1701) according to the manufacturer’s instructions. The resultant cDNA was analyzed by PCR with PowerPol 2 × PCR Mix (Abclonal, RK20719) using primers listed in supplementary table 5, and the PCR products were visualized by 2% agarose gel stained with Ethidium Bromide (Invitrogen, 15585011).

### Analysis of splicing intermediates

To measure the splicing intermediates, cDNA was generated from shCtrl- or shPTBP1-treated HeLa cells. Primers: F (targeting upstream constitutive exon), R (targeting downstream constitutive exon), Fa (targeting upstream intron near the 3’ splice site), Ra (targeting downstream intron near the 5’splice site), Fb (targeting downstream intron near the 3’ splice site), and Rb (targeting upstream intron near the 5’ splice site), were designed and used as previously described (Ule et al., 2006). For gel analysis, PCR was performed with 2 × M5 HiPer plus Taq HiFi PCR mix (Mei5bio, MF002 - plus-10) and the PCR products were fractionated on 8% TBE gel stained with SYBR Gold nucleic acid stain (Invitrogen, S11494). Quantitative PCR (qPCR) was performed with Hieff qPCR SYBR Green Master Mix (High Rox Plus) (Yeasen, 11203ES03) on a StepOnePlus real-time PCR machine (Applied Biosystems). Primers used are listed in supplementary table 5.

### Plasmids construction

Flag-tagged PTBP1 plasmid was generated as previously described (Xue et al., 2009). The HA-tagged PTBP1 was inserted into pcDNA3 between the *EcoR*I (NEB, R3101S) and *Not*I (NEB, R3189S) restriction sites. The PTBP1 synonymous mutant resistant to shRNA cleavage and C23S mutant for disrupting dimerization were generated following QuikChange site-directed mutagenesis protocol as previously described (Chen et al., 2018a). The plasmids for overexpressing the long and short isoforms of HERPUD2 were generated by cloning corresponding sequences into p3×Flag-CMV between the *EcoR*I and *BamH*I (NEB, R3136S) restriction sites, and the synonymous mutation of *HERPUD2* resistant to siRNAs was generated following QuikChange site-directed mutagenesis protocol as previously described (Chen et al., 2018a).

The *CTTN* minigene was described in a previous report (Xue et al., 2009). The *ZFYVE21* minigene (exon 5 to exon 7), and *HERPUD2* minigene (exon 6 to exon 8) , *TPM1* minigene (exon 5 to exon 7), and *TCFL5* minigene (exon 2 to exon 5) were constructed by inserting corresponding regions into pcDNA3 between *BamH*I and *EcoR*I restriction sites. Minigenes containing sQTLs were generated following QuikChange site-directed mutagenesis protocol as described (Chen et al., 2018a).

### Co-immunoprecipitation (Co-IP) and western blot

Co-IP and western blot were performed as previously described (Chen et al., 2018a). The anti-Flag (Sigma, F1804) and anti-HA (Sigma, H9658) antibodies were used for co-IP. The anti-PTBP1 (Abclonal, A18084), anti-DDDDK-tag (Abclonal, AE005), anti-HA-tag (Abclonal, AE008), and anti-β-Actin (Abclonal, AC026) antibodies were used for western blot.

### Protein purification, gel-shift assay, and SELEX

Protein purification, gel-shift assay, and SELEX were performed as previously described with minor modifications (Su et al., 2021). PTBP1-WT and PTBP1-C23S coding sequences were inserted into pET28a and transformed into Rosetta (DE3)-competent cells. The proteins were purified using the His GraviTrap column (GE Healthcare Life Sciences, 11-0033-99) by following the manufacture’s protocol. The Src-N1 3’ splice site RNA with 5’-Fam modification was synthesized by Sangon Biotech (Amir-Ahmady et al., 2005), and 0-350 nM proteins were incubated with 0.2 pmol RNA for the gel-shift experiments. Input RNA for SELEX was generated by *in vitro* transcription using the following dsDNA: GATCACTAATACGACTCACTATAGGGTACACGACGCTCTTCCGATCT (N10) AGATCGGAAGAGCACACGTCT (N represents the randomized nucleotides). The read count and enrichment (Z-score) for each heptamer were calculated as previously described (Su et al., 2021).

### RNA-tethering assay using CRISPR-dCasRx system

The sequences of PTBP1-WT and PTBP1-C23S were subcloned into pXR002 (EF1a-dCasRx-2A-EGFP 28 vector, Addgene, #109050) at restriction site *BsiW*I (NEB R0553S) using one-step cloning kit (Vazyme, C112). Oligos of single gRNAs and polycistronic gRNAs are synthesized by Sangon Biotech, annealed and inserted into pXR003 (gRNA cloning backbone vector, Addgene, #109053) between the two *Bpi*I (*Bbs*I) (Thermo Scientific, ER1011) restriction sites. PTBP1-dCasRx expressing plasmid and gRNA expressing plasmid were co-transfected into HeLa cells with GeneTwin transfection reagent (Biomed, TG101-02) according to the manufacturer’s instructions. Cells were harvested 72 h post-transfection for RT-PCR analysis.

### Doxycycline-inducible cell line construction

To create a Dox-inducible and pTRIPZ-mediated simultaneous knockdown and overexpression system, Flag-tagged PTBP1-WT and Flag-tagged PTBP1-C23S coding sequences resistant to shRNA cleavage were subcloned from pcDNA3 vectors into the pTRIPZ vector to replace RFP gene (Chen et al., 2018b; Paddison et al., 2004). The shRNA sequence targeting endogenous PTBP1 (5’-CGTGAAGATCCTGTTCAATA-3’) was inserted into the same pTRIPZ vector. To generate lentiviral particles, individual pTRIPZ-PTBP1 vectors were co-transfected with psPAX2 and pMD2.G into Lenti-X 293T cells (Takara, 632180), and the supernatant was collected twice at 48 h and 72 h post-transfection. To generate stable cell lines, HeLa cells were infected with lentiviral particle solution containing 8 μg/mL polybrene for 24 h and selected with puromycin (Solarbio, P8230) at a final concentration of 2 μg/ml 2 days after infection. Cells were treated with Dox (Clontech, 631311) at a final concentration of 1 μg/mL to induce the expression of exogenous PTBP1 and the transcription of shRNA to cleavage endogenous PTBP1 mRNAs. The Dox-containing medium was changed every 24 h to maintain the activity of the inducible promoter. To prevent the leaky expression in the absence of Dox, all the cells and lentivirus were prepared with TET Free Fetal Bovine Serum (Omega Scientific, FB-15).

### Cell proliferation and colony formation assays

Cell proliferation was performed with the MTS Kit (Promega, G3580) according to the manufacturer’s instructions. The dox-inducible HeLa cell lines were treated with Dox or an equal volume of solvent and seeded into 96-well plates at 1,000 cells per well. For validating the effects of *HERPUD2* isoforms, siRNAs (siNC or siHERPUD2 mix) were first transfected into HeLa cells with Lipofectamine RNAiMAX Reagent (Invitrogen, 13778150) according to the manufacturer’s instructions. After 24 h, vehicle or plasmids expressing different *HERPUD*2 isoforms were transfected into HeLa cells with GeneTwin transfection reagent. The cells were finally seeded into 96-well plates at 1,000 cells per well 24 h post-transfection. MTS reagent was directly added to the media, and the absorbance was measured at OD=490 nm after incubation at 37 °C for 1 h. For the colony formation assay, HeLa cell lines treated with Dox or an equal volume of solvent were seeded into a 6-well plate and maintained in DMEM containing 10% TET Free Fetal Bovine Serum for 10 days. Cells were washed with PBS twice, fixed with cold methanol, and stained with 0.1% crystal violet (Sigma, C3886) at RT for 15 min. The percentages of stained colonies were quantified by ImageJ software to determine the colony formation capacity.

### CRIC-seq data processing and analysis

Paired-end libraries of PTBP1 IP (+pCp), hnRNPA1 IP (+pCp), IgG IP (+pCp), PTBP1 IP (−pCp), and hnRNPA1 IP (−pCp) CRIC-seq were deep sequenced by Novogene. Raw data of all the CRIC-seq libraries were first aligned to the human genome as previously described (Liang Liang, 2022). To exclude the non-specific background, we developed a pipeline, as illustrated in Figure 1C, to identify high confidence chimeric reads. First, we separated the genome into 10-nt windows and combined the reads from IgG IP (+pCp) and PTBP1 IP (−pCp) CRIC-seq libraries as background. Then the chimeric reads of PTBP1 IP (+pCp) libraries will be retained for downstream analysis if they meet one of the following criteria: (1) their junctions sites and the junction sites of background chimeric reads are not present in the same pair of 10-nt windows; (2) their junction sites and the junction sites of background chimeric reads lie in the same pair of 10-nt windows, but the PTBP1 IP (+pCp) libraries showed 5-fold more chimeric reads linking the pair of 10-nt windows than the background. The retained PTBP1 IP (+pCp) chimeric reads owning junction sites within the same 10-nt windows were assembled into a cluster. The chimeras in the clusters supported by at least 2 reads were finally defined as high confidence reads representing authentic RNA-RNA interactions. The high confidence reads of hnRNPA1 IP (+pCp) were detected by the same pipeline.

### Genomic distribution of RBP-mediated high confidence chimeric reads

The genomic distribution of intramolecular high confidence chimeric reads was determined using gene annotation from the Gencode database (V19) and shown as pie graph (Figure S1K). The distribution of intramolecular high confidence chimeric reads around introns was shown in the heatmap (Figure S1L) generated by the pheatmap package (https://github.com/raivokolde/pheatmap) in R. To generate the heatmap, introns shorter than 100 kb and bound by PTBP1 (GSE42701) or hnRNPA1 (E-MTAB-3612) were selected (n=7,623, n=17,415). Then the selected introns were ranked in decreasing order of length. Next, 100-kb regions around the centers of introns were selected and separated into 1-kb bins. Finally, we calculated the density of intramolecular reads in every 1-kb bin and plotted them as a heatmap. The boundaries of the introns were depicted by a grey line.

### RBP motif enrichment analysis

The 193 RBP motifs were downloaded from the CISBP-RNA database (Homo sapiens, version 0.6) (Weirauch et al., 2014). The frequency of RBP motifs in the anchor regions without PTBP1 binding signals were counted by FIMO package (v5.0.4) (Grant et al., 2011). Then the motifs were ranked by the frequency in the anchor regions and the top3 motifs have shown as red dots. The protein-protein interaction (PPI) network of RBPs exhibited with top3 motifs were built by STRING program (v11.0, https://string-db.org/) (Mering et al., 2003) and displayed by the Cytoscape program (v3.8.2) (Shannon et al., 2003) with the default layout.

### 3D RNA map of PTBP1 mediated AS events

To build the 3D RNA map for PTBP1-regulated cassette exons (CE), we first identified significant CE events modulated by PTBP1 using the rMATS software (v4.0.2) based on the polyA+ RNA-seq data from the control and PTBP1 knockdown HeLa cells (GSE42701) (Xue et al., 2013). Significant CE events were filtered with the following criteria: (1) fragments per kilobase per million (FPKM) for the host gene was higher than 1; (2) the number of reads supporting exon inclusion and exclusion was more than 5; (3) the false discovery rate (FDR) was lower than 0.05; (4) the absolute value of difference of percent spliced in (|ΔPSI|=|PSI_WT_ - PSI_PTB KD_|) was greater than 0.1. Finally, we identified 215 significant CE events, including 80 PTBP1-enhanced and 135 PTBP1-repressed cases, among which 69 PTBP1-enhanced and 119 PTBP1-repressed cases own PTBP1 CRIC loops. Next, 2.5-kb regions (2 kb intronic regions and 500 nt exonic regions) around the exon-intron junction sites of the cassette exons and their flanking constitutive exons were selected from the 188 events containing CRIC loops, and then they were separated into 25 100-nt bins. We collected the high confidence chimeric reads located in these regions and normalized them by total RNA-seq data of HeLa cells. The normalized CRIC-seq read number was calculated as:

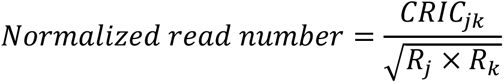

in which *CRIC_jk_* indicates the number of high confidence CRIC-seq chimeric reads linking the j^th^ and k^th^ bin, and *R_j_* and *R_k_* indicate the number of RNA-seq reads within the j^th^ and k^th^ 100-nt bins. Using the normalized chimeric reads, the normalized complexity map representing the 3D RNA map of PTBP1-regulated CE exons was calculated as previously described (Licatalosi et al., 2008; Xue et al., 2009). The 3D RNA map was displayed by combing every five 100-nt bins into a 500-nt bin. For comparison, a control 3D RNA map was generated using a set of 7,181 constitutive exons with the same calculation process as described above.

### Prediction of PTBP1-regulated splicing according to 3D RNA map

To predict cassette exons that may be regulated by PTBP1 based on the 3D RNA map, we first obtained 3,088 non-significantly changed CE events upon PTBP1 depletion with rMATS by setting FDR ≥ 0.05 and |ΔPSI| ≥ 0.1. Next, we calculated the normalized read number in 2.5-kb exon-intron junction regions of the potential CE events as described above. For a CE event, if ΔPSI≥0.1, and the normalized number of loops connecting the two ends of one intron was more than 4 and was 4-fold more than the loops bringing the two ends of the other intron and the loops straddling cassette exons, the CE event was predicted as a PTBP1-enhanced event. Conversely, if ΔPSI ≤ -0.1, and the normalized number of loops straddling cassette exons was more than 4 and was 4-fold more than the loops linking the two ends of both introns, the CE event was predicted as a PTBP1-repressed event. We finally obtained 329 potential PTBP1-enhanced and 64 PTBP1-repressed CE events.

### Cancer sQTLs and PPI network analysis

The sQTLs for 33 types of cancers were collected from the CancerSplicingQTL database (Tian et al., 2019). In total, we found 271,286 PTBP1 CRIC-seq clusters containing at least one sQTLs. To predict which CE event may be affected by these sQTLs, we first collected the PTBP1-regulated CE events (identified by RNA-seq and 3D RNA map) that had splice site-associated loops overlapped with sQTLs. Next, we used SELEX data to further filter the reserved CE events. The effect of sQTLs on PTBP1 binding affinity was evaluated by the fold change (FC) between heptamers containing reference sequence (reference heptamers) and heptamers containing sQTL (alternative heptamers). The CE events were retained if the |log_2_(FC)| is at least 0.6. As PTBP1 preferentially binds CU-rich elements, we focused on the sQTLs with CU-related transversion mutations (C/U to A/G, or A/G to C/U). Finally, we obtained 12 CE events of 11 genes containing cancer-related sQTLs. The protein-protein interaction (PPI) network for these 11 genes was generated by the STRING program (Mering et al., 2003) and displayed by the Cytoscape program (Shannon et al., 2003). Gene Ontology (GO) analysis was performed using the DAVID web server (v6.8) (Huang da et al., 2009a, b).

## FIGURES AND LEGENDS

**Figure S1.**
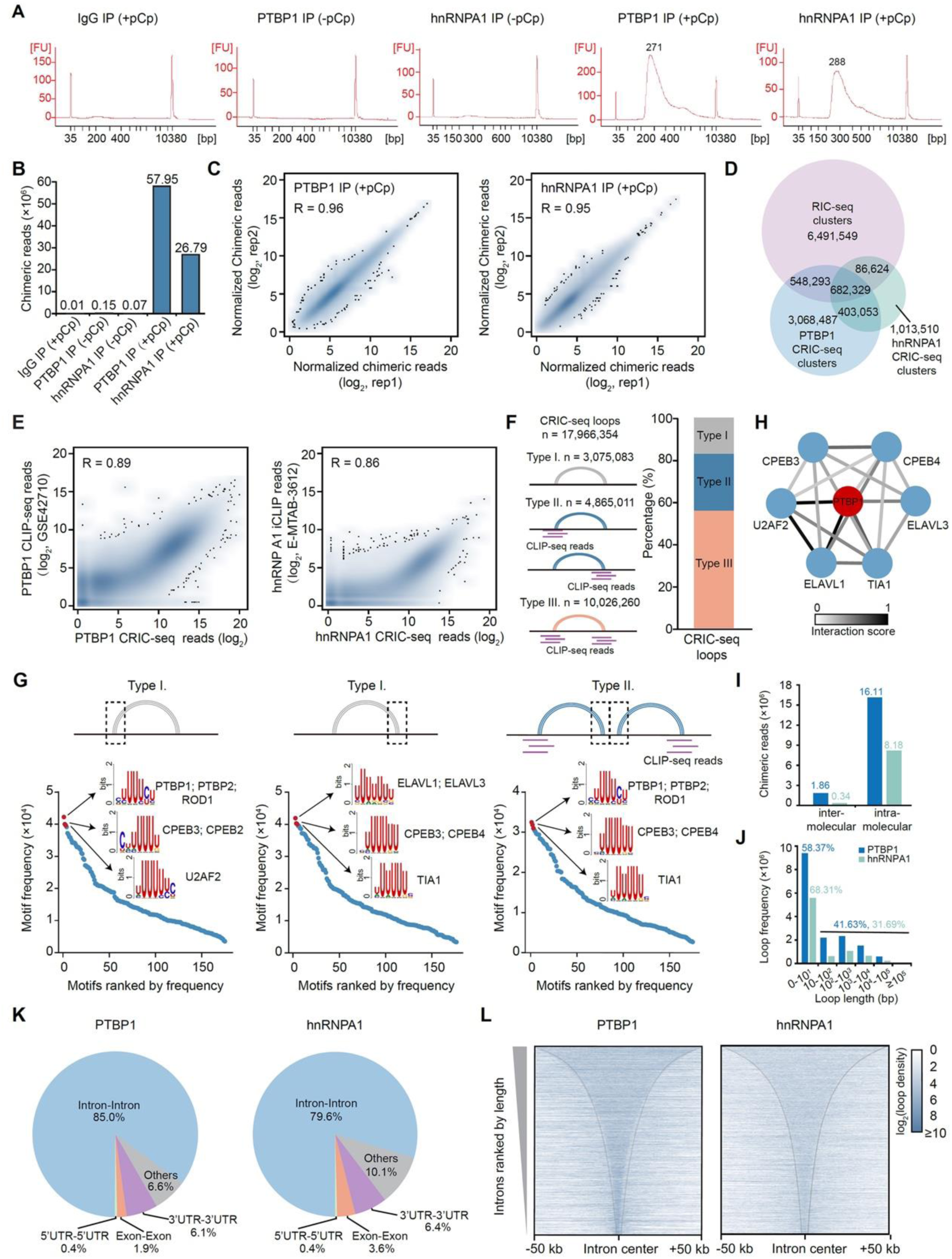
Validation of CRIC-seq method, related to Figure 1. (A) Size distribution of CRIC-seq libraries analyzed by the Agilent 2100 Bioanalyzer. (B) Chimeric read number for each CRIC-seq library. (C) Pearson’s correlation analysis shows a high correlation for PTBP1 and hnRNPA1 CRIC-seq replicates. (D) Venn diagram showing the overlap between RIC-seq, hnRNPA1 and PTBP1 CRIC-seq clusters. (E) PTBP1 CRIC-seq reads and CLIP-seq reads are highly correlated (*R*=0.89). hnRNPA1 CRIC-seq reads are also highly correlated with iCLIP reads (*R*=0.86). (F) CRIC-seq detected PTBP1 RNA loops could be classified into three types, and two of them have clear CLIP-seq signals. Type I loops do not have PTBP1 CLIP-seq signal, whereas CLIP-seq signals present at one arm or both arms in the other two types. (G) PTBP1 CRIC-seq reads without CLIP-seq binding evidence but contains CU- and U-rich motifs. (H) Protein-protein interaction network analysis showing PTBP1 potential interacting RBPs. Color of the edge indicates interaction scores from the STRING database. (I) Bar graph showing the numbers of high confidence inter- and intramolecular chimeric reads detected by PTBP1 CRIC-seq. (J) The distribution of spanning distance for intramolecular RNA-RNA interactions. (K) RNA-RNA interaction landscape for intramolecular PTBP1 and hnRNPA1 CRIC-seq chimeric reads. (L) The aggregation plot shows that PTBP1 and hnRNPA1-mediated intramolecular loops are largely confined within introns. Introns are ranked based on their length. The gray line indicates the boundary of introns. Color intensity indicates CRIC-seq read density.

**Figure S2.**
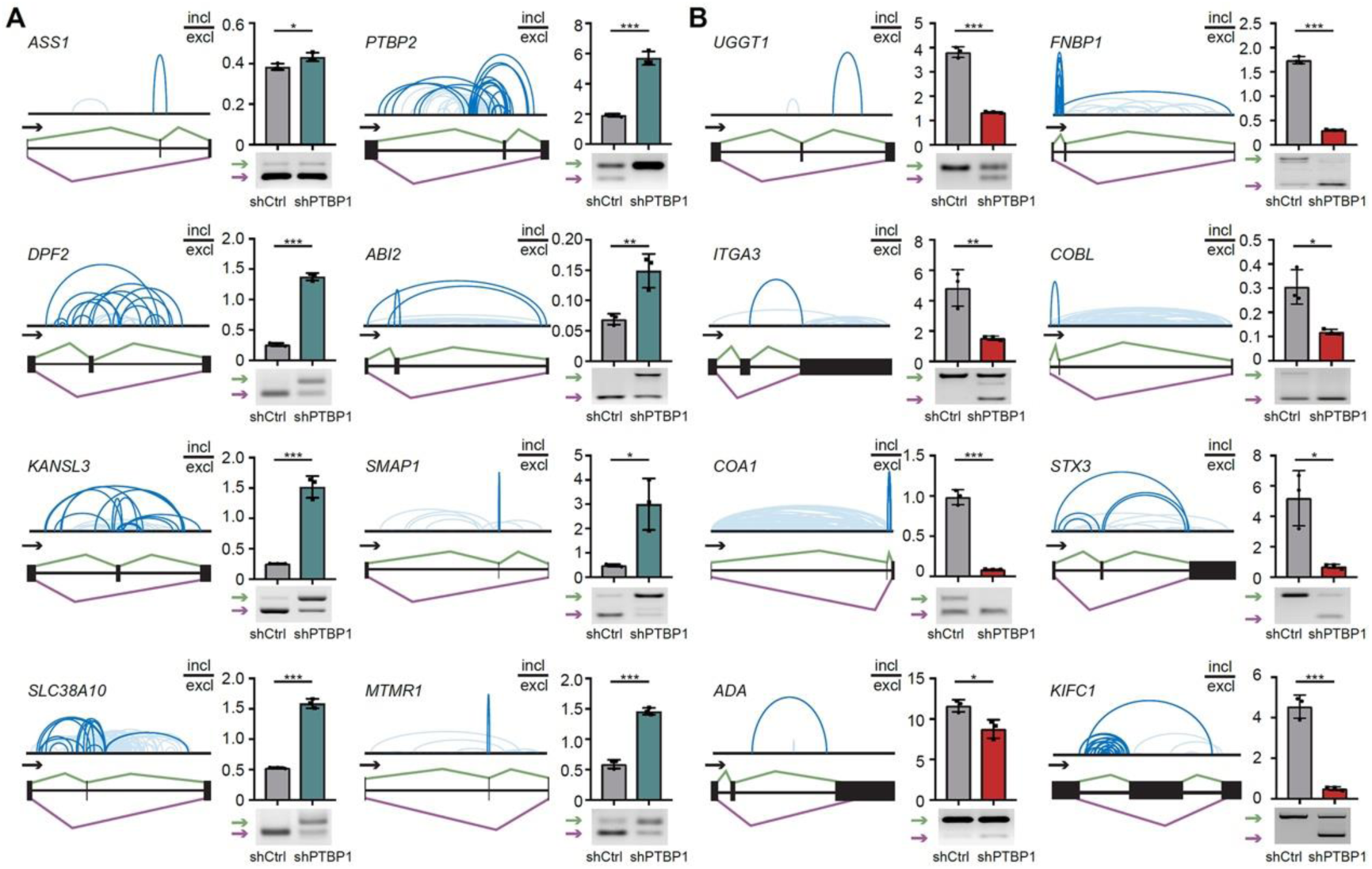
Validation of PTBP1 CRIC-seq-detected splicing events, related to Figure 2. (A) Validation of PTBP1-repressed cassette exons by RT-PCR. (B) Validation of PTBP1-activated cassette exons by RT-PCR. Data in (A) and (B) are mean ± s.d.; n=3 biological replicates, two-tailed, unpaired *t*-test. ^∗^*P* < 0.05, ^∗∗^*P* < 0.01, ^∗∗∗^*P* < 0.001.

**Figure S3.**
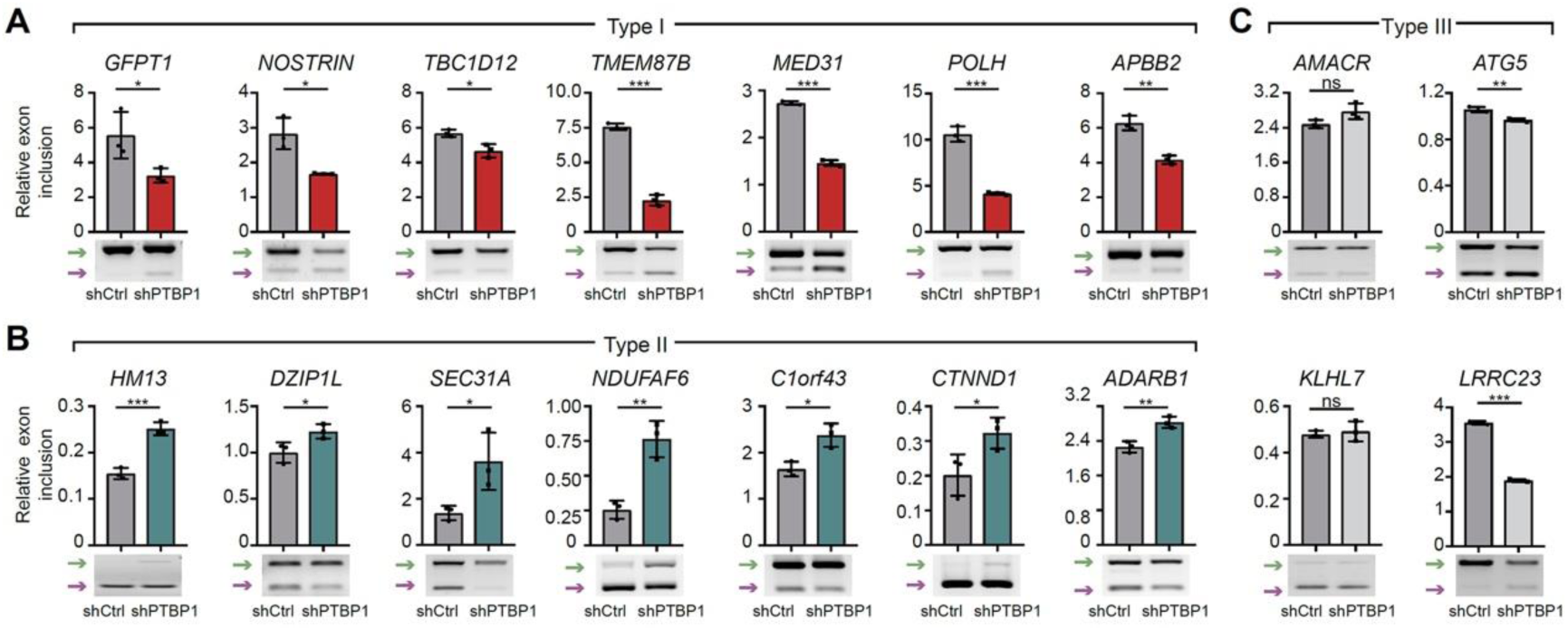
Validation of predicted splicing events based on 3D RNA map, related to Figure 3. Validation of cassette exons predicted to be positively (A) or negatively (B) regulated by PTBP1. *AMACR*, *KLHL7*, *ATG5*, and *LRRC23* cassette exons not regulated by PTBP1 served as negative controls (C). Data in (A), (B), and (C) are mean ± s.d.; n=3 biological replicates, two-tailed, unpaired *t*-test. ^ns^*P* >0.05, ^∗^*P* < 0.05, ^∗∗^*P* < 0.01, ^∗∗∗^*P* < 0.001.

**Figure S4.**
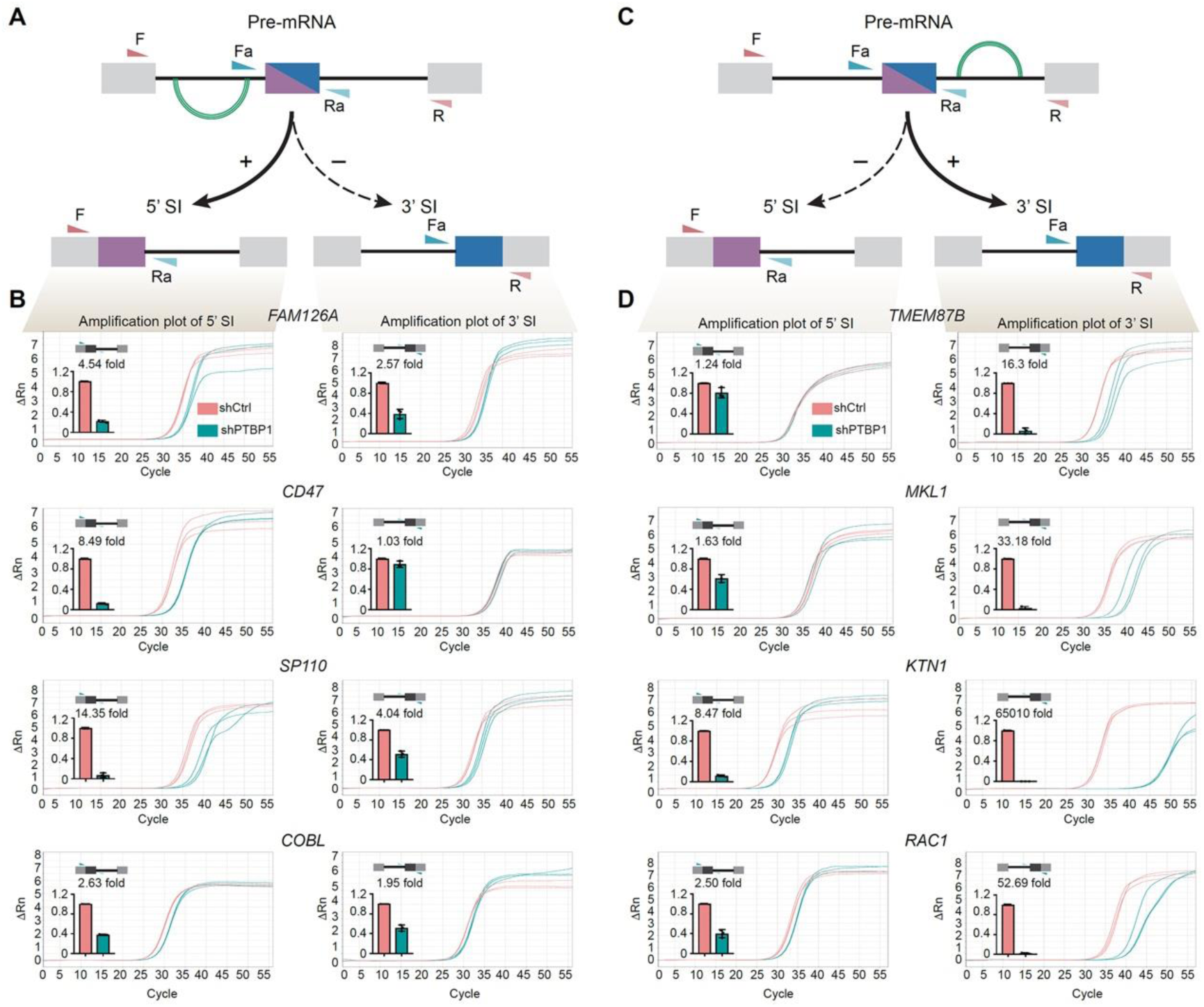
RT-qPCR to quantify splicing intermediates, related to Figure 4. (A) Schematic diagram of the quantification of splicing intermediates for PTBP1-mediated RNA loops in upstream introns by RT-qPCR. PTBP1-mediated RNA loops in upstream introns are shown in green arc lines. The intronic loops will promote the removal of upstream introns, leading to the significantly increased production of 5’ splicing intermediates. (B) RT-qPCR to quantify the splicing intermediates of three pre-mRNAs (*FAM126A*, *CD47*, *SP110*, and *COBL*) which contain more loops in the upstream intron. (C) Diagram of the quantification of splicing intermediates for PTBP1-mediated RNA loops in downstream introns by RT-qPCR. (D) RT-qPCR to quantify splicing intermediates of four pre-mRNAs (*TMEM87B*, *MKL1*, *KTN1,* and *RAC1*) which contain more loops in the downstream intron. Data in (B) and (D) are mean ± s.d.; n=3 biological replicates, two-tailed, unpaired *t*-test.

**Figure S5.**
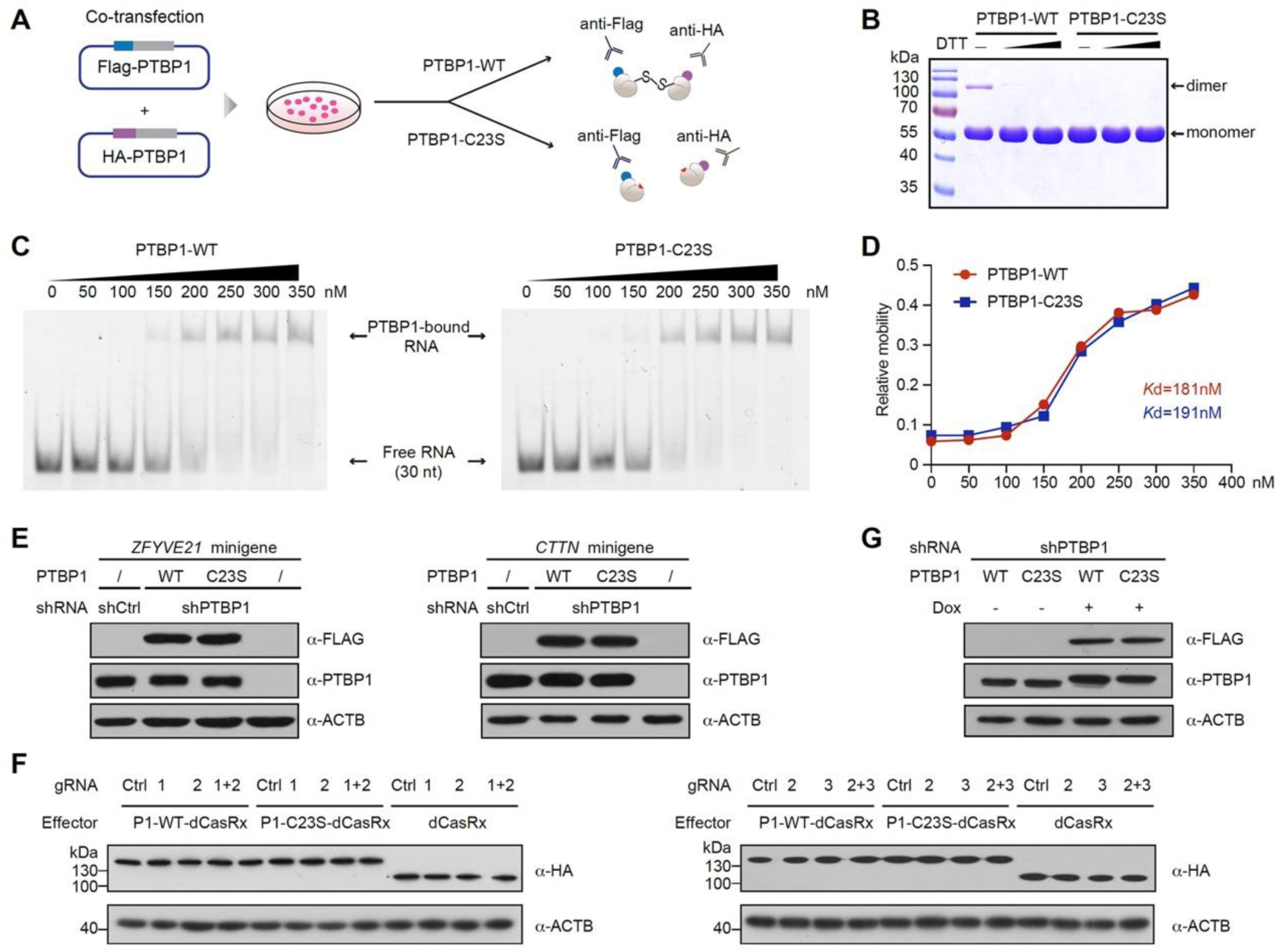
C23S is required for PTBP1 dimerization but not for its binding to RNA, related to Figure 5. (A) Diagram of reciprocal co-immunoprecipitation with Flag- and HA-tagged wild-type (WT) and C23S mutant proteins. (B) SDS-PAGE shows the dimerization of purified PTBP1-WT proteins rather than PTBP1-C23S. DTT, dithiothreitol. (C) The gel-shift assay shows that PTBP1-WT and PTBP1-C23S have a comparable ability to bind CU-rich RNA. (D) The relative mobility of PTBP1-WT and C23S proteins is similar. Data are plotted with the mean value from two biological replicates. (E) Western blot showing comparable expression of exogenously PTBP1-WT and PTBP1-C23S in shPTBP1-treated HeLa cells compared with shCtrl-treated cells. (F) Western blot showing similar expression levels of exogenously PTBP1-dCasRx and dCasRx. P1, PTBP1; gRNA, guide RNA. (G) Western blot shows Dox inducible expression of PTBP1-WT or C23S mutants in shPTBP1-treated HeLa cells.

**Figure S6.**
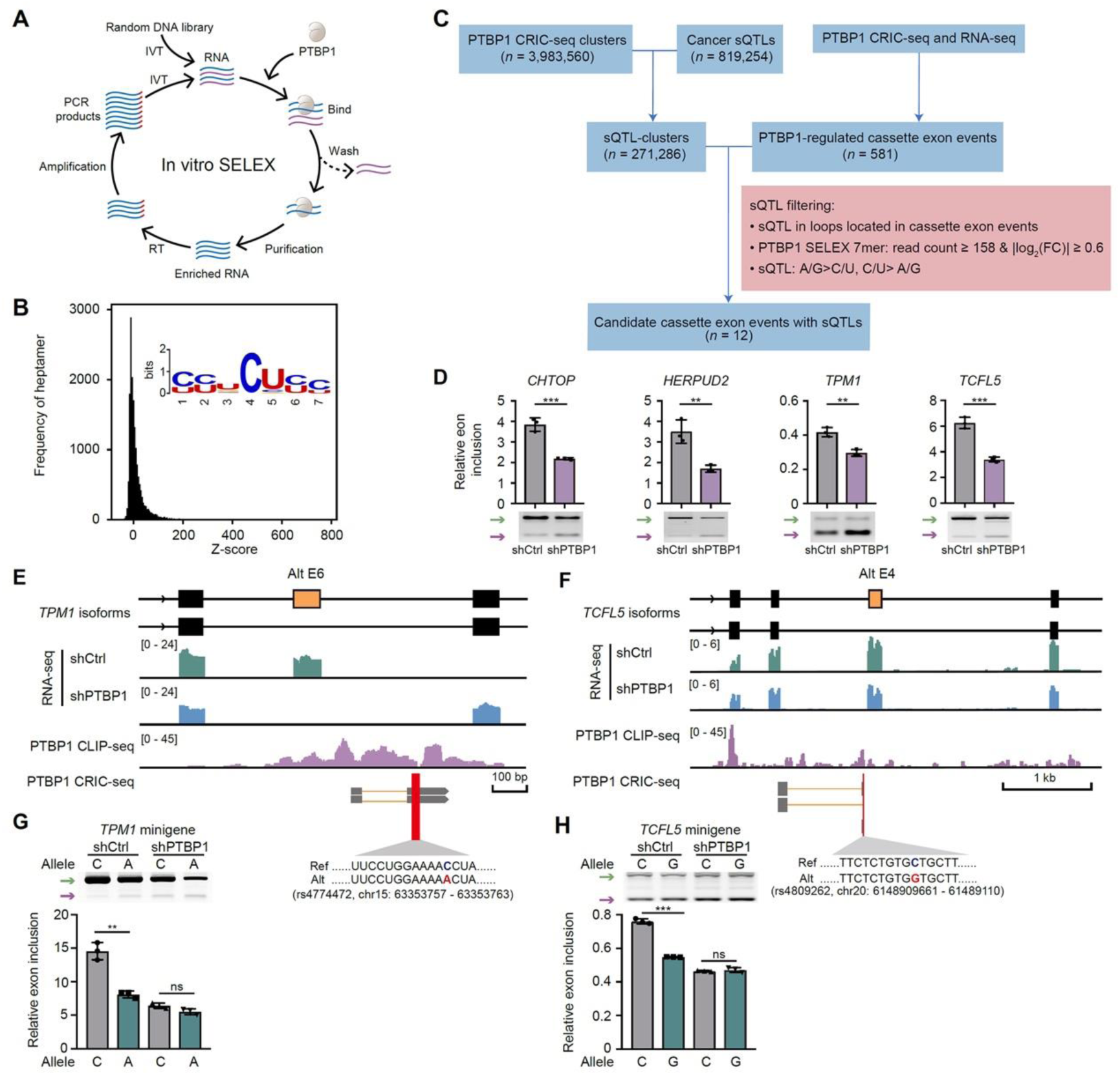
Validation of sQTLs on splicing misregulation, related to Figure 6. (A) Schematic diagram showing the *in vitro* SELEX strategy to identify RNA sequences preferentially bound by PTBP1. (B) Histogram showing the frequency of RNA heptamers enriched by the PTBP1 SELEX assay. (C) The analysis pipeline to identify cassette exon splicing events that contain sQTLs and the mutations may affect splicing by changing the RNA binding affinity of PTBP1. (D) Validation of cassette exon splicing by semiquantitative RT-PCR. Green arrows indicate the long isoforms, and purple arrows indicate the short isoforms. (E, F) Snapshot of *TPM1* exon 6 and *TCFL5* exon 4 and their flanking regions. RNA-seq signals in shCtrl and shPTBP1 HeLa cells are shown in green and blue. CLIP-seq reads are shown in purple. CRIC-seq clusters with the sQTL are shown in gray boxes. (G, H) The splicing of sQTLs containing *TPM1* and *TCFL5* reporters showed decreased exon usage in shCtrl-treated HeLa cells, and the mutants failed to respond to PTBP1 knockdown. Green arrows indicate the long isoforms, and purple arrows indicate the short isoforms. Data are mean ± s.d.; n=3 biological replicates, two-tailed, unpaired *t*-test. ^ns^*P* >0.05, ^∗∗^*P* < 0.01, ^∗∗∗^*P* < 0.001.

